# Restraint Validation of Biomolecular Structures Determined by NMR in the Protein Data Bank

**DOI:** 10.1101/2024.01.15.575520

**Authors:** Kumaran Baskaran, Eliza Ploskon, Roberto Tejero, Masashi Yokochi, Deborah Harrus, Yuhe Liang, Ezra Peisach, Irina Persikova, Theresa A. Ramelot, Monica Sekharan, James Tolchard, John D. Westbrook, Benjamin Bardiaux, Charles D. Schwieters, Ardan Patwardhan, Sameer Velankar, Stephen K. Burley, Genji Kurisu, Jeffrey C. Hoch, Gaetano T. Montelione, Geerten W. Vuister, Jasmine Y. Young

## Abstract

Biomolecular structure analysis from experimental NMR studies generally relies on restraints derived from a combination of experimental and knowledge-based data. A challenge for the structural biology community has been a lack of standards for representing these restraints, preventing the establishment of uniform methods of model-vs-data structure validation against restraints and limiting interoperability between restraint-based structure modeling programs. The NMR exchange (NEF) and NMR-STAR formats provide a standardized approach for representing commonly used NMR restraints. Using these restraint formats, a standardized validation system for assessing structural models of biopolymers against restraints has been developed and implemented in the wwPDB OneDep data deposition-validation-biocuration system. The resulting wwPDB Restraint Violation Report provides a model vs. data assessment of biomolecule structures determined using distance and dihedral restraints, with extensions to other restraint types currently being implemented. These tools are useful for assessing NMR models, as well as for assessing biomolecular structure predictions based on distance restraints.

**Graphical Abstract:** 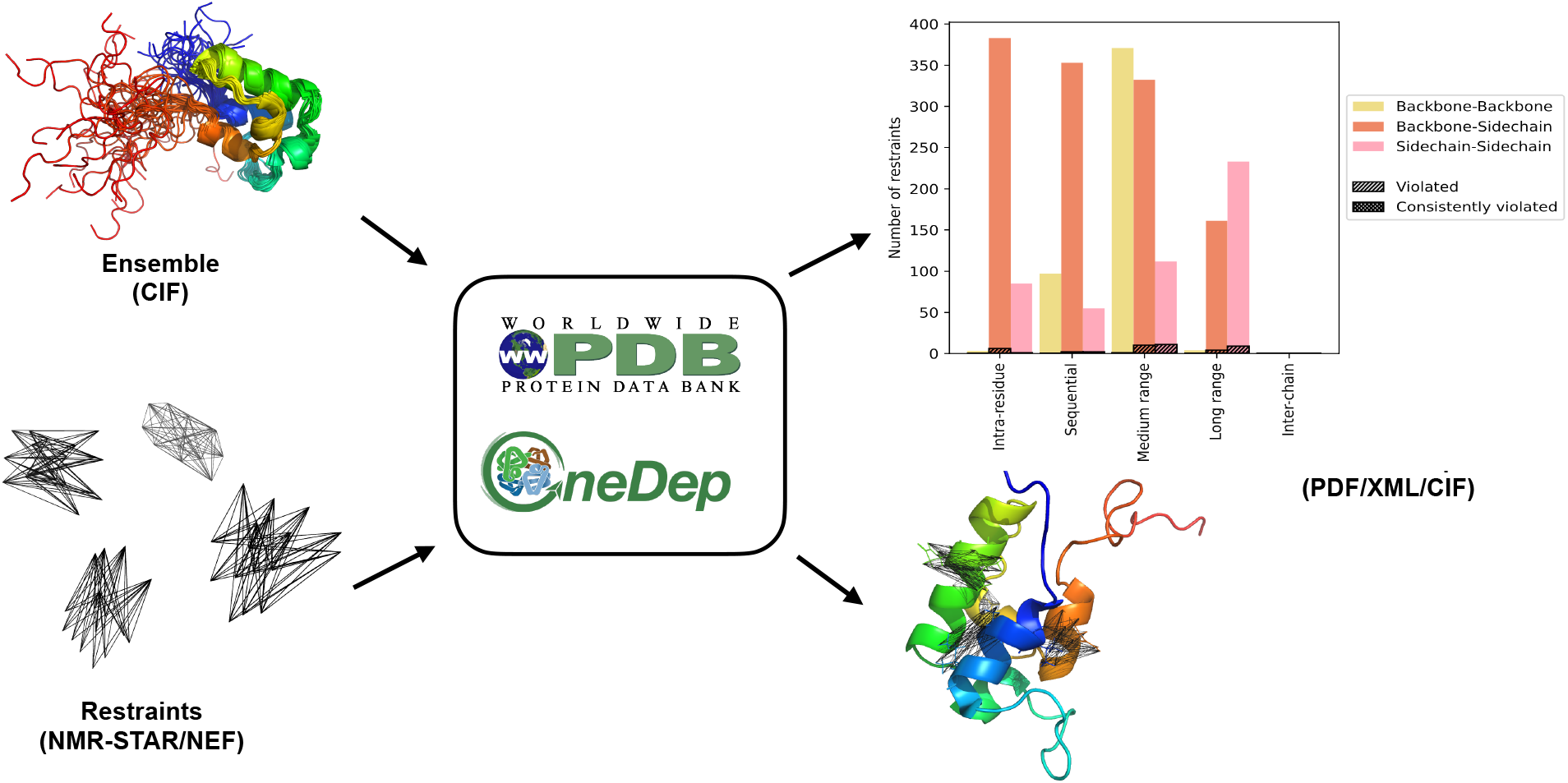

**Highlights:**

- PDB Structure Validation Report expanded to include Restraint Analysis
- NMR Exchange Format (NEF) and NMR-STAR for distance restraint representation
- Standard distance and dihedral restraint formats for model vs. restraint assessment
- Standardized restraint formats provide interoperability between modeling programs

## INTRODUCTION

### Structure determination using NMR

Nuclear Magnetic Resonance (NMR) spectroscopy is a versatile experimental technique used not only for structure determination but also to probe conformational dynamics and interactions of biomolecules. NMR-derived biomolecular structures are primarily modeled using estimates of interatomic distances and dihedral angles between atoms or groups of atoms in the form of distance and dihedral angle restraints. These restraints are provided as input to restrained molecular dynamics or other structural modeling programs, which incorporate covalent bond geometry and conformational energy force fields, and output a collection of atomic-resolution models, the so-called “NMR ensemble”, in which each conformer is a best fit to all of the experimental restraints. Note that this is a looser definition than the concept of a “thermodynamic statistical ensemble” in statistical mechanics, which describes the Boltzmann distribution of the conformations contributing to ensemble-averaged measurable parameters.

The Protein Data Bank (PDB) is in its 52^nd^ year of continuous operation. Established in 1971 as the first open-access digital data resource in biology (Protein_Data_Bank, 1971), it currently houses > 200,000 experimentally determined 3D structures of proteins and nucleic acids (DNA and RNA) and their complexes with one another and with small-molecule ligands (*e.g*., enzyme cofactors, inhibitors, peptides, and drugs). The Worldwide Protein Data Bank (wwpdb.org) partnership currently includes five full members [i.e., Research Collaboratory on Structural Bioinformatics Protein Data Bank (RCSB PDB), Protein Data Bank in Europe (PDBe), Protein Data Bank Japan (PDBj), Biological Magnetic Resonance Bank (BMRB), and Electron Microscopy Data Bank EMDB] and one associate member [the Protein Data Bank China

(PDBc)], which jointly manage the PDB, EMDB, and BMRB core archives (Berman *et al*., 2003; wwPDB Consortium, 2019). All PDB data are made available by wwPDB partners under the most permissive Creative Commons CC0 license. wwPDB members are committed to ensuring that structural biology data are FAIR (Findable, Accessible, Interoperable, and Reusable) (Wilkinson *et al*., 2016) and FACT (Fairness, Accuracy, Confidentiality, and Transparency) (van der Aalst *et al*., 2017). Objective assessment and validation of biomolecular structure models based on NMR, X-ray crystallography, cryogenic electron microscopy, small angle X-ray scattering, and integrative structural biology methods are important and ongoing activities of the wwPDB (Berman *et al*., 2003; Read *et al*., 2011; Henderson *et al*., 2012; Montelione *et al*., 2013; Trewhella *et al*., 2013; Sali *et al*., 2015).

Biomolecular structure validation includes two general classes of assessment, knowledge-based validation, in which the model(s) are assessed in light of what is known about biomolecular structure from the existing database of experimental structures, and model vs. data validation, in which consistency is assessed between the structural model(s) and experimental data obtained for the subject biomolecule (Montelione *et al*., 2013; Rosato *et al*., 2013). The latter is crucially dependent upon the description of the measured quantities (*e.g*., nuclear Overhauser effects - NOEs, residual dipolar couplings – RDCs, etc.) as a function of the atomic coordinates. Accurate experimental models should score well across the multiple metrics available for these two assessment categories (Bhattacharya *et al*., 2007; Rosato *et al*., 2013). In such assessment methods, it is also important to estimate and consider the uncertainty of the model. This is generally done by comparing models generated from multiple runs of the model generation software to identify regions that are consistently modeled (*i.e*., the “well-defined” regions of the model) and those that are not consistently modeled from the available data and methods (*i.e*., the “not-well-defined” regions of the model) (Hyberts *et al*., 1992; Snyder and Montelione, 2005; Kirchner and Güntert, 2011; Montelione *et al*., 2013; Rosato *et al*., 2013; Snyder *et al*., 2014). More rigorously, the precision of the model could be estimated by the propagation of experimental uncertainties using Bayesian methods (Rieping *et al*., 2005), but so far, this has been done in only a small number of biomolecular structure studies. Consensus recommendations for tools useful for knowledge-based validation and conventions for defining “well-defined regions” of biomolecular structure models have been provided by the wwPDB NMR Structure Validation Task Force (Montelione *et al*., 2013) and implemented in the wwPDB NMR Validation Report (Gore *et al*., 2017) using standardized knowledge-based validation methods (Chen *et al*., 2010; Rosato *et al*., 2013; Vuister *et al*., 2014), with the understanding that NMR structure model validation methods continue to evolve and improve.

Ideally, experimental structures should be validated against the primary experimental data. Although several methods have been developed and are in use for validating structures against NOE, chemical shift, RDC, or other experimental data, no consensus has yet emerged on best practices for collecting, archiving, and using these data for structure validation. As most biomolecular NMR structures are determined using distance and dihedral angle restraints derived from such primary data, a minimal criterion for NMR structure validation is the assessment of the deposited models against these derived restraint data. In assessing models against these restraints, one of the most challenging issues is that different structure generation software tools utilize NMR-derived distance restraints in different ways and formats. While tools have been developed to convert between some NMR restraint formats (Vranken *et al*., 2005; Tejero *et al*., 2013; CCPN, 2023), and some large-scale remediation efforts have been performed at the Biological Magnetic Resonance Data Bank (Nederveen *et al*., 2005), in some cases it is challenging to represent distance-restraint information used by one structure generation program accurately in the restraint functions of a different program without the active involvement of the software developers in this process. These challenges have been addressed, at least in part, through the development of the NMR Exchange Format (NEF, (Gutmanas *et al*., 2015)), designed to provide reliable interoperability between NMR software programs and structure generation programs in particular. Its design was strengthened by involving software developers in creating and testing all aspects, including the NMR restraint representations.

Here, we report an accurate two-way interconversion between the NEF restraint format and the NMR-STAR restraint format which is the NMR data archive format of the wwPDB. The current NEF convertor supports the translation of chemical shifts, distance, and dihedral angle restraints necessary for the validation process from NEF version 1.1 to NMR-STAR version 3.2. This NEF / NMR-STAR converter is available through wwPDB GitHub repository (https://github.com/wwPDB/py-wwpdb_utils_nmr). We use it to implement a new restraint validation component of the wwPDB NMR Structure Validation Report, which builds on distance and dihedral-angle restraint representations in the NMR-STAR format, generating both a human-readable report in PDF format and machine-readable format in CIF (Westbrook *et al*., 2022). It also provides an XML representation of the data for further computer analysis. These innovations have been validated against the stand-alone software PDBStat for restraint format interconversion (Tejero *et al*., 2013). They have also been validated against the CcpNmr Analysis version-3 program suite (Skinner *et al*., 2016) and NEF implementations in the NMR structure calculation programs Xplor-NIH (Schwieters *et al*., 2006) and ARIA (Rieping *et al*., 2007), with implementation in the program CYANA/CANDID (Güntert and Buchner, 2015) close to completion as well. Together, these tools allow for straightforward generation and validation of experimental NMR-derived structure models against distance and dihedral-angle restraints using restraints in either NEF or NMR-STAR format.

In this paper, we describe the development of a comprehensive NMR model *versus* distance and dihedral-angle restraint data validation report for the wwPDB. Future extensions to other NMR restraint types (*e.g*., RDCs) can be readily implemented within the same framework. It is anticipated that this model vs. data NMR Restraint Validation Report, together with the already available knowledge-based NMR Structure Validation Report, will provide a more comprehensive and objective assessment of the reliability of biomolecular structures determined by NMR methods.

## RESULTS

### Restraint validation

Checking the validity of a given restraint between two atoms on a given model is not as trivial as it might appear, as several complicating aspects must be considered for a proper analysis.

Typically, although not in all programs, the distance restraint is modeled using a cost function corresponding to a square well potential and defined by the lower limit and upper limit of the distance between the two atoms. If the measured distance in the model falls between these bounds, then the restraint is not violated, otherwise, it is deemed violated. For methods that do not use bounds, e.g. in the log-harmonic potential as implemented in ARIA (Nilges *et al*., 2008), the respective software is expected to specify how the strength of potentials needs to be translated into equivalent lower and upper bounds.

The complications for a meaningful restraint violation analysis arise from several factors. First, the overlap of resonances and ambiguity in their assignments need to be accounted for. Often, a so-called “r^-6^ sum” over all the distances contributing to the restraint can be used to address such overlap and ambiguity (Hyberts *et al*., 1992; Nilges, 1995; Bassolino-Klimas *et al*., 1996). Second, all the restraints derived from the NMR experiments result from the spatial and temporal averages of the underlying dynamics exhibited by the biomolecule, and not all restraints may be fully satisfied at all instances of time. Molecular dynamics simulations, such as those employed to calculate structures based on NMR data, ensure that on average, all restraints are satisfied for a maximum amount of time by minimizing the total energy of the system. A small fraction of restraints may be violated at every instance, but the set of restraints violated in each instance is different so that no restraint is consistently violated. Third, NMR structure calculations typically result in an ensemble or collection of conformers. This conformational multiplicity results from both the experimental uncertainty and potentially from actual conformational variability in the sample. The NMR-VTF has recommended that the NMR ensemble should be analyzed in terms of either well-defined or not-well-defined regions, unless it is explicitly modeled as a Boltzmann conformational distrubution (Montelione *et al*., 2013), reflecting the fact, by definition, that the biomolecular conformation in not-well-defined regions is not reliably modeled. Unusual dihedral angle values and steric clashes in the latter regions are not considered to be significant. Additionally, the extent to which dynamical information can be faithfully represented in a fixed-size ensemble is an unanswered question.

Highly similar considerations apply to the validation of all types of NMR-derived restraints. Whereas assignment ambiguity is generally not pertinent for dihedral-angle restraints, conformational averaging is a similarly complicating factor. RDC restraints are also conformationally averaged, albeit on different timescales compared to the NOE-derived distance restraint. Additionally, analysis of RDC satisfaction requires establishing the alignment tensor (Losonczi *et al*., 1999), with its own associated uncertainties.

### Types of restraints and their validation

The distance restraint between two atoms is a derived quantity, sometimes originating from more than one spectrum or more than one peak in a single spectrum (e.g., from symmetric NOESY peaks). In a conventional NMR structure determination workflow, the chemical shift assignments are first derived from one or more (heteronuclear) through-bond type experiments, which are then used to assign the peaks of through-space NOESY-type spectra. The volumes or intensities of these assigned NOESY peaks are then used to estimate the distance (or upper-bound distance) between pairs of atoms. It is not uncommon to have identical chemical shifts within the spectral resolution for more than one atom. For example, the three protons of a methyl group usually have degenerate chemical shifts, and chemical shifts of different methyl groups from the same or different residues may also overlap. Such overlapping chemical shifts lead to ambiguous chemical shift assignments, which results in ambiguous restraints. Below, we discuss the typical cases a distance restraint validation procedure needs to accommodate.

### Type 1: Unambiguous distance restraints

Unambiguous distance restraints can often be derived from well-resolved NOESY peaks between two atoms. The chemical shifts of the atoms involved in this type of restraint are non-degenerate and unambiguously assigned. Let *r_ij_* be the distance between atom *i* and *j* in the molecular model (Figure 1A), *d_min_ (i,j)* is the lower bound and *d_max_(i,j)* is the upper bound of the distance restraint. If *d_min_ (i,j)* ≤ *r_ij_* ≤ *d_max_(i,j),* the restraint is not violated, otherwise the violation is calculated as the lowest of the | r_ij_ - d_min_ (i,j) | or | r_ij_ - d_max_ (i,j)|.

**Fig. 1.**
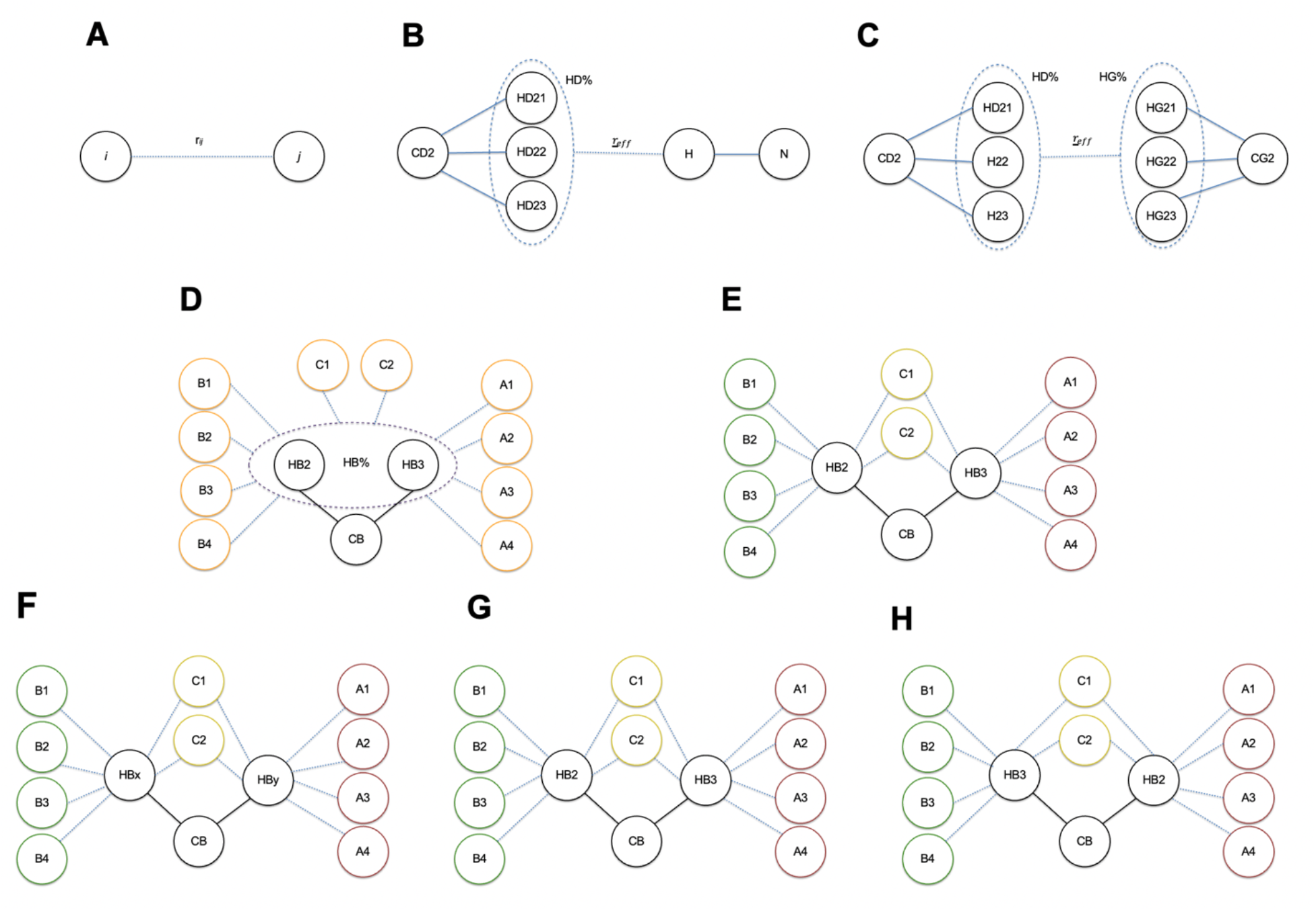
(A) Type1: Distance restraint between atoms i and j. (B) Type 2: Distance restraint between an atom and a group of atoms (C) Type 2: Distance restraint between two groups of atoms. (D) Type 3: Distance restraint between non-stereo specifically assigned atoms with degenerate chemical shifts and groups of atoms. (E) Type3: Distance restraint between stereo- specifically assigned atoms with non-degenerate chemical shifts and group of atoms. (F) Type3: Distance restraint between non-stereo specifically assigned atoms with non-degenerate chemical shifts and groups of atoms. (G,H) possible assignments for (F).

### Type 2 Ambiguous restraints involving resonances with degenerate chemical shifts

Degenerate chemical shifts (*e.g*., those of magnetically equivalent methyl protons) give rise to ambiguous distance restraints. The NEF standard provides for the wild-card “%” identifier, *e.g*., HB%, to allow for ambiguous restraints involving such degenerate chemical shifts. To a first approximation, the NOESY peak between the degenerate resonance, such as a methyl group, and another atom will have NOE contributions from each of the contributing protons (Figure 1(b), (c)), which varies inversely to the distance. As an approximation, an effective distance (*r*_*eff*_) can be calculated using the r^-6^ sum (Eq. 1) of the pairwise distances between all pairs of atoms contributing to the NOE peak (Nilges, 1995), and any violation of the restraint is assessed using the resulting *r*_*eff*_. *If d_min_ (i,j)* ≤ *r*_*eff*_ ≤ *d_max_(i,j),* then the restraint is not violated, otherwise the violation can be calculated as min (| *r*_*eff*_- d_min_ (i,j) |, | *r*_*eff*_- d_max_ (i,j)|), where *r*_*eff*_ is given by

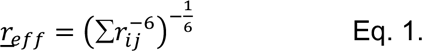

The r^-6^ sum distance restraint, *r*_*eff*_, has the feature that it is dominated by the shortest distance to the set of ambiguously assigned (or degenerate) atoms. It can be used not only for degenerate methyl or methylene protons but also for any two or more protons involving degenerate resonances.

### Type 3: Restraints between atoms involving resonances of stereo-specifically- or individually-assignable atoms

Prochiral methylene protons, individually-assignable amide NH_2_ groups (for example, from asparagine and glutamine), and aromatic ring protons (for example, from phenylalanine and tyrosine side chains), may or may not have degenerate chemical shifts. If they are non-degenerate, they can potentially be stereo-specifically or individually assigned. Isopropyl methyl groups (for example, from valine and leucine) are also prochiral and generally admit stereo- specific assignments. However, unless individual or stereospecific assignments are established using specific experimental or computational methods, these groups of resonances are also treated using ambiguous restraints. This ambiguity can be addressed by effectively treating the two separate resonances as though they are degenerate, and summing the volumes (or intensities) to create an ambiguous r*^-6^* sum restraint (Nilges, 1995; Tejero *et al*., 2013). Although ambiguous restraints may be defined differently in input to different structure generation programs, the current wwPDB Restraint Validation Report validates the model against ambiguous restraints, assuming an r^-6^ sum interpretation (Eq 1, above). NEF includes a robust standard to handle these situations, linking the NMR resonance assignment intimately with the calculated structures in the NMR ensemble (see Methods).

#### Case 3.1: Degenerate chemical shifts (HB% case, r^-6^ sum)

If a group of protons, such as those of methylene groups (or other degenerate proton resonance groups), have degenerate chemical shifts, then they can be treated like Type 2 restraints described above, and the r^-6^ sum method (Eq. 1) is used to validate the model against the restraint (Figure 1D).

#### Case 3.2: Non-degenerate and stereo-specifically assigned

If the chemical shifts of a group of protons are non-degenerate and if they are stereo-specifically (or individually) assigned, then these restraints are treated as either a Type 1 or Type 2 restraint depending on whether the other atom is a single atom or group of atoms, respectively (Figure 1E).

#### Case 3.3: Non-degenerate and ambiguously assigned

If the chemical shifts of a group of protons are non-degenerate and if they are not stereo- specifically assigned, the situation is much more complex as the various structure generation algorithms employ different approaches in dealing with this issue. Thus, crucial information needs to be captured and adequately handled in the restraint validation protocols, which traditionally has presented a serious problem. The issue is best illustrated with an example. Let us assume that the chemical shifts of a group of protons, *e.g*., two methylene protons, are non-degenerate but cannot be assigned stereo-specifically. Following the NEF standard, these protons should be labeled with the “x” and “y” identifiers, *i.e*., HBx and HBy for a pair of methylene protons attached to CB. Assume that a set of NOESY peak-derived distance restraints was observed for HBx to atoms B1, B2, B3, B4, C1 and C2 and another set of restraints observed for HBy to atoms A1, A2, A3, A4, C1 and C2 (Figure 1F). The example indicates that in addition to the set of common restraints, *i.e*., to C1 and C2, there are also restraints exclusive to either HBx or HBy. It is *a priori* undefined whether the stereo-specific HB2 atom in the molecular structure maps onto HBx, and the HB3 atom to HBy, or *vice versa*.

One straightforward (though imperfect) approach in dealing with this issue is to collapse the two restraints to resonances HBx and HBy into one restraint involving an ambiguous HB%, with some form of treatment of the restraint limits, *e.g*., on the basis of the originating peak intensities or by taking either the shortest (most restricting) or longest (least restricting) limit. In one simple but common implementation, the two stereo-specifically distinct resonances are treated as degenerate, their intensities are summed, and a r^-6^ sum restraint (Eq. 1) is created (Nilges, 1995; Tejero *et al*., 2013). In this approach, a Case 3.3 HBx/HBy restraint has been converted to a Case 3.1 ambiguous restraint. The current wwPDB Restraint Validation Report validates the model against restraints involving nondegenerate and ambiguously assigned proton groups, assuming an r^-6^ sum interpretation (Eq 1, above).

In some methods for addressing ambiguous stereochemical assignments, the two prochiral atoms are represented by a pseudoatom at the midpoint, and a single restraint is made to this pseudoatom ensuring that, considering the ambiguity of the stereospecific assignment, the longest proton-proton distance satisfies the restraint (Wüthrich *et al*., 1983; Fletcher *et al*., 1996). This pseudoatom restraint will be looser than the r^-6^ restraint outlined above. If the pseudoatom upperbound distance restraint reported in the restraint file satisfies the r_eff_ upper- bound distance calculated from the r^-6^ sum approach, it will also satisfy the upper-bound distance to the pseudoatom. Accordingly, the current wwPDB Restraint Validation Report validates the model against pseudoatom restraints, assuming an r^-6^ sum interpretation (Eq 1, above).

Another approach, implemented by structure calculation programs like ARIA, Xplor-NIH or Cyana, is the concept of “floating chirality” (Folmer *et al*., 1997). In the course of the structure calculation, the program adopts the most favorable mapping for all pairs at any time, thus minimizing the resulting restraint energy. At some point during the calculation, typically in light of sufficient consistency, the mapping can be fixed. However, such structure-based stereo-specific assignments cannot usually be obtained for all pairs of prochiral or individually-assignable atoms. In the past, structure calculation programs like ARIA/CYANA (Brünger *et al*., 1998) have provided such mapping information for subsets of restraints in the form of the so-called “float- files” or “stereo.aco” files, respectively. Unfortunately, this information has mostly been lost during the deposition process using data supplied in a program-specific format, and this approach leads to inconsistency between atoms defined in the restraints and the model files.

Extensive efforts to re-capture these mappings required great efforts and were only partially successful (Doreleijers *et al*., 2012). Chemical shift prediction from model structures could offer a path to resolving ambiguous stereo-specific assignments (Weiss and Hoch, 1987).

By documenting the actual restraints used in the structure calculation in NEF format and reporting structural data in PDBx / mmCIF format, this issue is solved. Using NEF, the stereo- specific mapping, *i.e*., HB2 onto HBx and HB3 onto HBy or vice versa, can and should be documented for each atomic position using the ambiguity tag, <u>atom_site.pdbx_atom_ambiguity</u>, and the restraint is validated accordingly (Figure 1G,H). Note that this mapping could differ for each model of the structural ensemble. In the absence of such a documented mapping using a NEF-PDBx / mmCIF pair, and to avoid the introduction of an erroneous restraint validation assessment, the restraint validation algorithm interprets both HBx and HBy as HB%, effectively treating them as Case 3.1 degenerate restraints, as described above. This procedure ensures that violations are not wrongly reported and that appropriate restraint and restraint-violation counts are maintained.

The usage of the NEF-PDBx / mmCIF pair and ambiguity tag has the added advantage of solving a long-standing problem involving restraints involving slowly-rotating phenylalanine or tyrosine aromatic side chain protons, *i.e*., in the case of non-degenerate HD1/HD2 and HE1/HE2 chemical shifts. Structurally, the HD1/HD2 and HE1/HE2 designations, as well as the CD1/CD2 and CE1/CE2 designations, are defined by the χ2 dihedral angle, and a small rotation beyond 180° will structurally swap CD1/HD1/CE1/HE1 with CD2/HD2/CE2/HE2. This, however, could have detrimental consequences for restraints formulated in terms of HD1/HD2/HE1/HE2 as large errors would be introduced by such a swap. This can be resolved using the HDx/HDy/HEx/HEy NEF nomenclature and ambiguity tag mapping in the PDBx / mmCIF file, as the mapping effectively provides the correct atomic coordinates to be used for the restraint validation. Note that restraints formulated in terms of the wild-card HD% or HE% atoms are always evaluated correctly.

Going forward, the resolution of Case 3 non-degenerate ambiguously assigned restraints depends on the generation of consistent pairs of NEF-PDBx / mmCIF and atomic coordinate files. Given the involvement of the software development community in the NEF project, we are confident that the common NMR structure calculation programs are poised to create such pairs, thereby assuring the correct data interpretation and a correct restraint validation.

### Dihedral Angle Restraints

A dihedral angle is defined as the angle between half-planes defined by two sets of three atoms having two atoms in common. Customarily, the common atoms are bonded, and each bonded to the other defining atoms. In proteins, the backbone dihedral angles ɸ and Ѱ, defined by the backbone atoms C-N-CA-C and N-CA-C-N, respectively, provide crucial information to describe the main-chain geometry. The Ramachandran plot, which visualizes the distribution of these two backbone dihedral angles on a ɸ-Ѱ graph, is widely used by various structure validation program suites (Laskowski *et al*., 1993; Lovell *et al*., 2003; Chen *et al*., 2010) to assess the quality of the structure.

For proteins, dihedral-angle restraints can be derived using backbone chemical shift data (Cheung *et al*., 2010; Shen and Bax, 2010; 2015). In the NEF format, any dihedral-angle restraint is defined using its four relevant atoms, a target value, upper-bound and lower-bound values. The sign of these angles indicates whether the angle is measured counterclockwise or clockwise. The convenient use of positive and negative signs in representing angles makes the upper and lower bounds an arbitrary choice. Without the target value, it is hard to tell which side of the angular region between the upper and lower bound is the allowed region. Figure 2 illustrates how different choices of target values render either the acute or the obtuse angle as allowed regions.

**Fig. 2.**
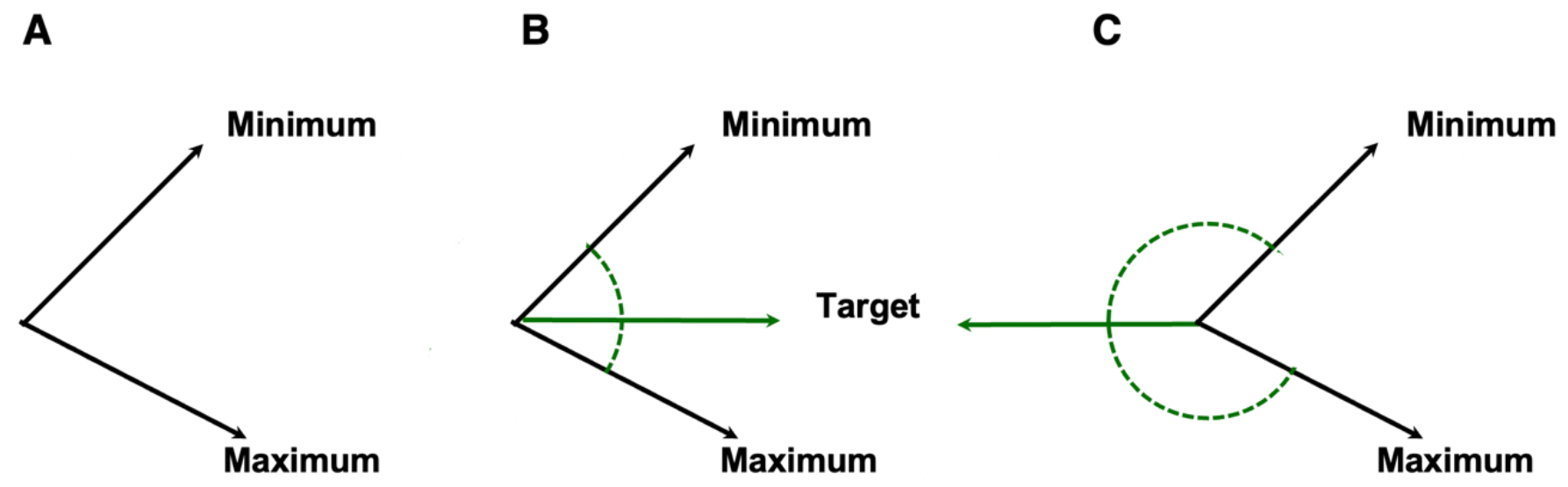
(A) Angle restraint without target value. (B,C) Two possible target values for a given set of minimum and maximum, which makes either the counterclockwise (B) or the clockwise(C) angular region the allowed region.

For example, let ϕ*_mesaured_* be the measured backbone dihedral angle in a given model. If both ϕ*_target_* and ϕ*_mesaured_* are in the angular region between ϕ*_min_* and ϕ*_max_* then the dihedral restraint is not violated; otherwise, it is violated, and the violation ϕ*_violation_* is calculated as

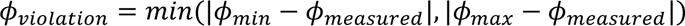

Dihedral angle restraints may also be ambiguous, *i.e*., they may define multiple, discontinuous regions of the ϕ − ψ map. The target value may potentially be assigned to more than one value to define multiple conformations that are consistent with the data. Ambiguous dihedral restraints are defined in NEF as a set of restraints using the combination of _nef_dihedral_restraint.restraint_id and _nef_dihedral_restraint.restraint_combination_id. The ambiguous restraint is considered violated only if all of the possible restraints in the set are violated.

### wwPDB OneDep deposition and validation

The global wwPDB OneDep tool (Young *et al*., 2017) supports deposition, validation (Gore *et al*., 2017; Feng *et al*., 2021), and biocuration (Young *et al*., 2018) for macromolecular structures determined by macromolecular crystallography (MX), 3D electron microscopy (3DEM), and NMR since its launch in 2014. Recently, the OneDep has been enhanced to further support NMR restraint data generated by community software in either NEF (V1.1) (Gutmanas *et al*., 2015) or NMR-STAR (V3.2) (Ulrich *et al*., 2019) format. The OneDep deposition interface allows authors to upload a single combined NMR data file that includes required chemical shift and restraint data and optional peak list data in either NEF or NMR-STAR format while (presently) continuing to support native file formats from community software (*e.g*., CYANA, CNS, and Xplor-NIH). The latter option, however, will be phased out in the near future, in consultation with the NMR community, as many problems associated with reliable interpretation are now addressed by using the NEF-PDBx / mmCIF pair of NMR and structural data (*vide infra*). We encourage software developers to generate biomolecule structure deposition data in NMR-STAR / NEF formats for NMR data and PDBx / mmCIF format for atomic coordinate files for proper validation.

Assigned chemical shifts and the experimental restraint data are mandatory for deposition to the PDB of a biomolecular structure solved using NMR spectroscopy. Depositors are also highly encouraged to provide NOESY peak lists, as well as other relevant NMR information, such as RDC data, as part of their NEF file. The validation workflow converts the uploaded NEF data into NMR-STAR format, the archival format of NMR data in the PDB and BMRB Core Archives. The validation report is generated using the NMR data in NMR-STAR format and coordinate data in PDBx/mmCIF format.

The NMR-STAR file is used for restraints because this is the archival format of the BMRB. The NEF file is more lightweight, and better suited as an interoperable exchange format. As part of this project, NMR-STAR -> NEF and NEF -> NMR-STAR converters have been developed (https://github.com/wwPDB/py-wwpdb_utils_nmr). Notably, the NEF allows for documenting the cross-database entry identifiers to allow for related information to be retrieved and analyzed.

The NMR-STAR/NEF NMR data file should contain the following mandatory data, encoded as so-called blocks/save frames in accordance with the respective defined data formats:

1. Sequence information
2. Assigned chemical shift data
3. Restraint data (various types)

Depositors are encouraged to also provide as much metadata as possible in their NMR-STAR/NEF file using the tags defined by their respective data dictionaries. However, some necessary metadata will be collected through the wwPDB deposition user interface and added to the NMR-STAR or NEF data file. Upon file upload, OneDep provides the following diagnostics:

1. Identifies the file type as an NMR unified data.
2. Validates the NMR data, including checking NMR data content and providing data diagnostics such as identifying unusual chemical shift data values, i.e, chemical shift values outside of the expected range. The various checks provide warnings for depositors to review (Table 1).
3. Cross-checks the sequence between the atomic coordinate and the chemical shift files and provides a sequence alignment for depositors to review.

**Table 1.**
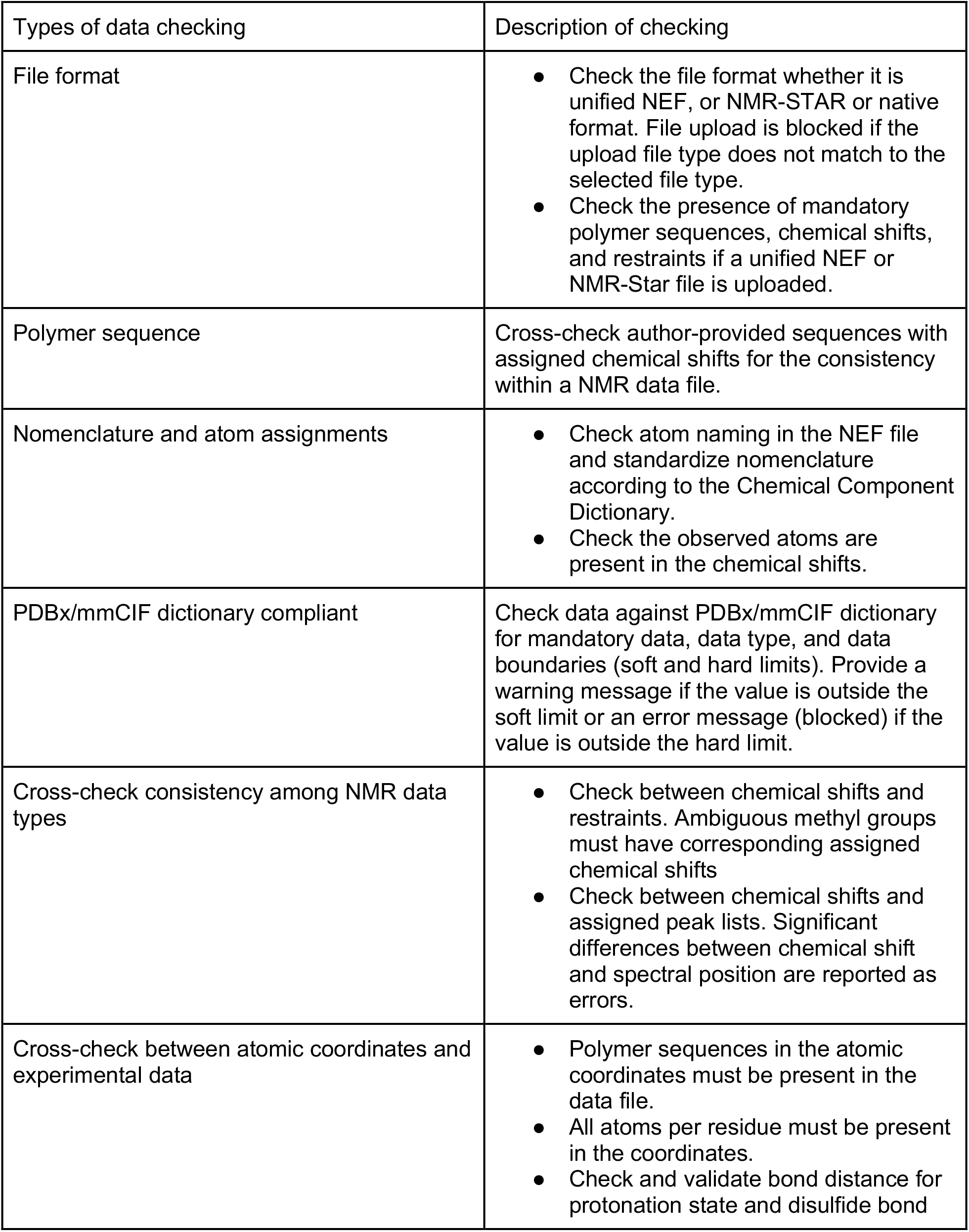
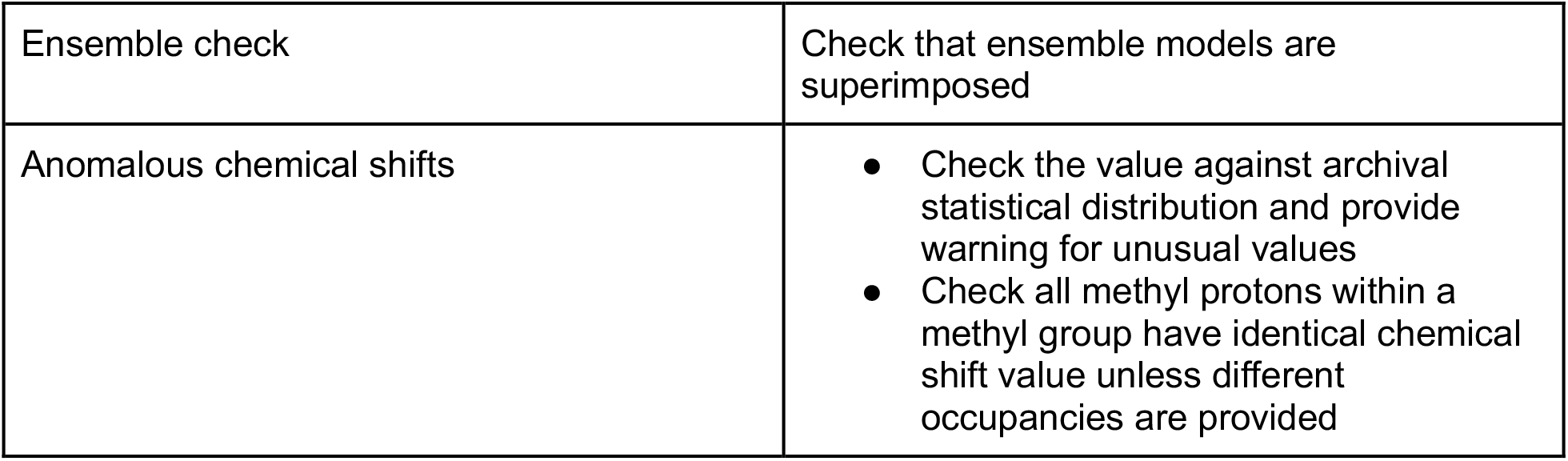
NMR data checking at OneDep deposition.

| Types of data checking | Description of checking |
| --- | --- |
| File format | • Check the file format whether it is unified NEF, or NMR-STAR or native format. File upload is blocked if the upload file type does not match to the selected file type.<br>• Check the presence of mandatory polymer sequences, chemical shifts, and restraints if a unified NEF or NMR-Star file is uploaded. |
| Polymer sequence | Cross-check author-provided sequences with assigned chemical shifts for the consistency within a NMR data file. |
| Nomenclature and atom assignments | • Check atom naming in the NEF file and standardize nomenclature according to the Chemical Component Dictionary.<br>• Check the observed atoms are present in the chemical shifts. |
| PDBx/mmCIF dictionary compliant | Check data against PDBx/mmCIF dictionary for mandatory data, data type, and data boundaries (soft and hard limits). Provide a warning message if the value is outside the soft limit or an error message (blocked) if the value is outside the hard limit. |
| Cross-check consistency among NMR data types | • Check between chemical shifts and restraints. Ambiguous methyl groups must have corresponding assigned chemical shifts<br>• Check between chemical shifts and assigned peak lists. Significant differences between chemical shift and spectral position are reported as errors. |
| Cross-check between atomic coordinates and experimental data | • Polymer sequences in the atomic coordinates must be present in the data file.<br>• All atoms per residue must be present in the coordinates.<br>• Check and validate bond distance for protonation state and disulfide bond |
| Ensemble check | Check that ensemble models are superimposed |
| Anomalous chemical shifts | • Check the value against archival statistical distribution and provide warning for unusual values<br>• Check all methyl protons within a methyl group have identical chemical shift value unless different occupancies are provided |

Once the uploaded NMR data file has passed the file check, some metadata are automatically parsed at the deposition interface for authors to review and are also captured in the atomic coordinate file to reference the corresponding NMR data. If a unified NMR data file is uploaded, it is then passed to the wwPDB validation package for generation of the wwPDB Validation Report (Gore *et al*., 2017; Feng *et al*., 2021), which includes restraint validation as outlined here. The resulting preliminary NMR Structure Validation Report is provided at the deposition interface for the author’s review and correction of their data as needed. Subsequently, the official wwPDB NMR Structure Validation Report is generated by the wwPDB biocurators after data processing and is sent back to the authors for the journal manuscript review process. The wwPDB Validation Reports are provided in PDF, PDBx/mmCIF, and XML formats. At the time of release for PDB entries, the wwPDB validation reports are generated for public distribution at https://ftp.wwpdb.org/pub/pdb/validation_reports and the unified NMR data are made available at https://ftp.wwpdb.org/pub/pdb/data/structures/divided/nmr_data/ in both NEF and NMR-STAR formats.

### Restraint violation analysis in wwPDB Validation Reports

The distance and dihedral-angle restraint analysis is presented in the wwPDB Validation Report in sections 8, 9, and 10; an example is provided as Supplementary Material. Section 8 describes the overall summary of the deposited restraints of all categories and restraint violations in different bins. For distance restraints, a grouped data classification is widespread in the NMR community and was recommended by the wwPDB NMR Validation Task Force (Montelione *et al*., 2013). It provides a simple and convenient overview of the available data and their agreement with the molecular structure.

The conformationally restricting restraints are counted in categories of intra-residue, sequential, medium range (|i-j| > 1 and |i-j| < 5), long-range (|i-j| ≥ 5), and inter-chain restraints, and listed (Table 2) along with the number of hydrogen bond restraints and disulfide bond restraints. Table 2 shows a summary of restraints data for a representative NMR structure, PDB ID 7M5T (Anishchenko *et al*., 2021), as an example. The full validation report is available on the PDB entry summary page. Restraints involving interatomic distances that are already restrained by covalent structure are considered as *redundant restraints,* and multiple restraints between the same two atoms (e.g., restraints between atoms A to B and B to A, or two different distance restraints between the same atom pair A - B) are considered to be *duplicate restraints*. When duplicate restraints have different upper bounds, the looser restraint is used in the assessment. Distance restraint values that do not restrict the conformations of the intervening dihedral angles are also identified, and these *non-conformationally-restricting restraints* are excluded from the restraint validation analysis. Additional details of this restraint filtering are presented in Tejero et al., 2013. If an atom involved in a restraint has no corresponding atom in the coordinate file, it is counted as an *unmapped restraint*. Duplicate, redundant, non-conformationally-restricting, and unmapped restraints are reported back to the user and excluded from the reported statistics.

**Table 2.** Conformationally-restricting restraints for PDB ID 7m5t.

| Description | Value |
| --- | --- |
| Total distance restraints | 2189 |
| Intra-residue ($\vert i-j\vert =0$) | 471 |
| Sequential ($\vert i-j\vert =1$) | 505 |
| Medium range ($\vert i-j\vert >1$ and $\vert i-j\vert <5$) | 675 |
| Long range ($\vert i-j\vert \geq 5$) | 398 |
| Inter-chain | 0 |
| Hydrogen bond restraints | 140 |
| Disulfide bond restraints | 0 |
| Total dihedral-angle restraints | 175 |
| Number of unmapped restraints | 0 |
| Number of restraints per residue | 23.6 |
| Number of long-range restraints per residue | 4.0 |

All conformationally-restricting restraints are validated against structural results from each model in the NMR ensemble. If the measured distance between a pair of atoms (or the r_eff_ for ambiguous restraints computed as the r^-6^ sum distance) in a given model lies between the upper and the lower bound of the corresponding distance restraint as described above, then the restraint is not violated. If the measured distance in a model lies outside the boundaries defined by the restraint, then the absolute difference between the measured value and the nearest boundary is reported as the violation value. The results are reported, and binned into small, medium, and large violation categories based on the magnitude of the violation values. In each bin, the average number of violations per model is calculated by dividing the total number of violations in each bin by the size of the ensemble. The maximum value of the violation in each bin is also reported. Table 3 lists distance violations per bin in the *de novo* designed protein PDB ID 7M5T (Anishchenko *et al*., 2021) as an example. If dihedral-angle restraints were included, similar overall and violation statistics are also provided for these. Violations less than 0.1 Å for distance restraints and less than 1° for angle restraints, which may have come from round-off errors, are excluded from the statistics.

**Table 3.** Average number of distance violations per model using PDB ID 7M5T as an example.

| Bins | Average number of violations per model | Max (Å) |
| --- | --- | --- |
| 0.1 - 0.2 Å (Small) | 3.6 | 0.2 |
| 0.2 - 0.5 Å (Medium) | 0.9 | 0.32 |
| > 0.5 Å (Large) | None | None |

Sections 9 and 10 in the Validation Report provide a detailed analysis of distance and dihedral-angle restraints. Both sections have similar subsections and contents for their respective restraints categories and hence are discussed here together. Sections 9.1 and 10.1 describe the summary of violations in different restraint categories. For each category, the table provides; the total number of restraints, the percentage with respect to the total, the number of violated restraints, and the percentage with respect to both that particular category and to the total number of restraints. Restraints that are violated in at least one model are counted as *violated*, and restraints that are violated in all the models are counted as *consistently violated*. The information in the table is also provided as a bar chart, which gives a straightforward overview of any consistent violations. The example for PDB ID 7M5T (Figure 3) shows a typical pattern, with only a few violated restraints and no consistently violated restraints.

**Fig. 3.**
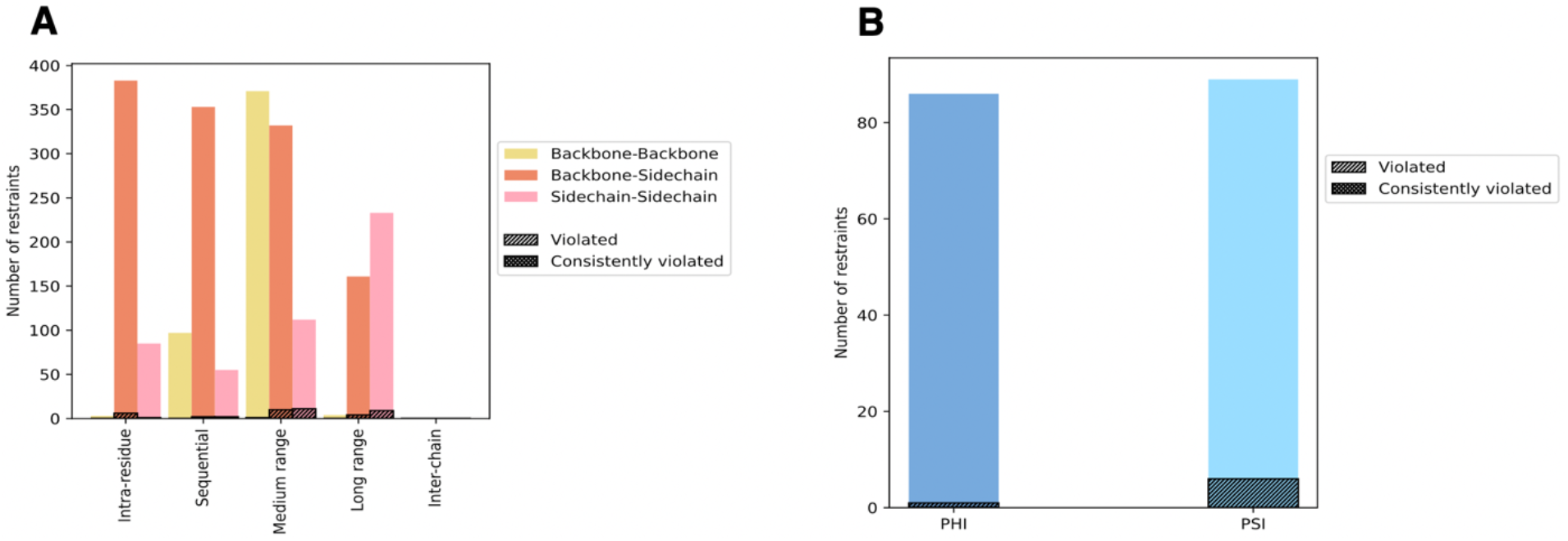
Bar graph distribution of (A) distance and (B) dihedral angle restraints of PDB ID 7M5T. Violated and consistently violated portions are shown in different hash patterns.

The following subsections in the report, sections 9.2 and 10.2, provide the violation statistics for each model. The number of violations in each model and the mean, median, standard deviation, and maximum values are listed in a table and are also presented as a bar chart in the report.

Figure 4 shows the per-model bar chart for PDB ID 7M5T (Anishchenko *et al*., 2021). The total number of violations for each model (∼ 6) is low. The distribution of violations, as indicated by the blue bars and indicators for median and mean (∼ 0.17 Å) distance restraint violation, is also low and highly similar for all models, suggesting that no model represents an outlier. Models 13, 14, and 20, however, appear slightly better for the agreement of local conformations with the experimental data, as these models show no violations in the intra-residue and sequential categories.

**Fig. 4.**
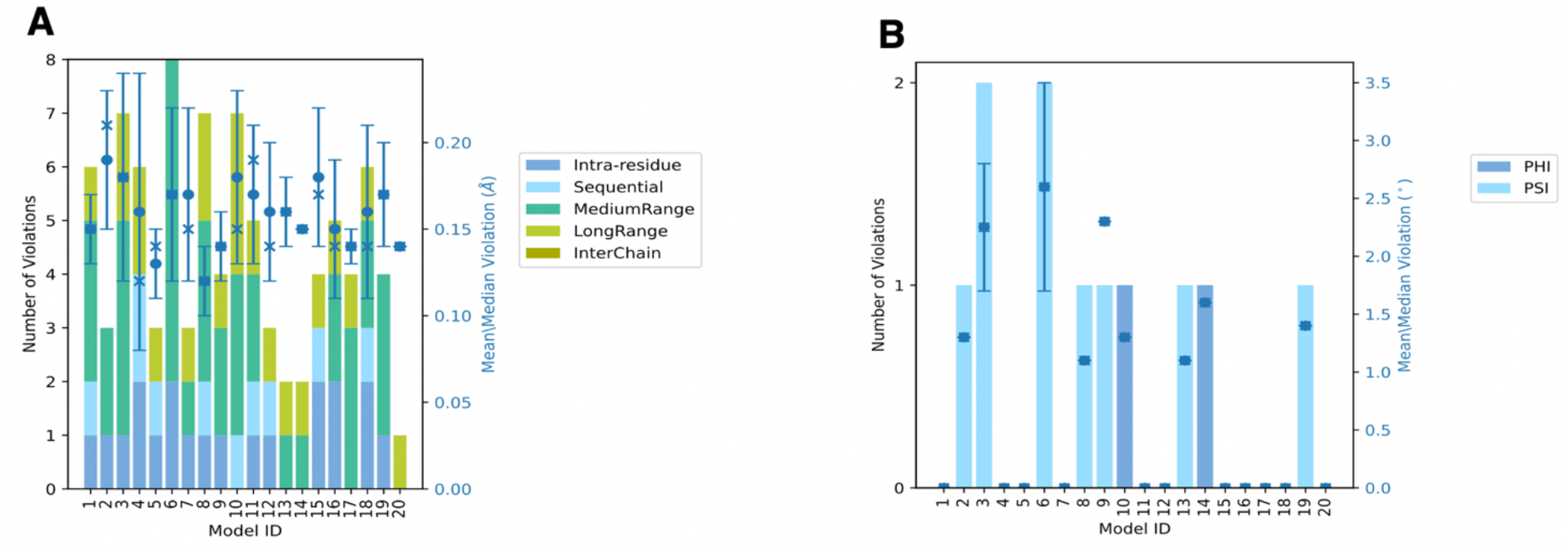
Per-model (A) distance and (B) dihedral violation statistics of PDB ID 7M5T. The mean (dot), median (x), and the standard deviation (error bar) of the violation are shown in blue with respect to the y axis on the right.

The distance and dihedral angle violation statistics for the ensemble are presented in sections 9.3 and 10.3 of the report, respectively. The table in these sections lists the number of violations for a given fraction of the ensemble. The number of restraints violated in all models, *i.e*., the consistently violated restraints, are also listed. The bar chart (Figure 5) shows the violation statistics for the ensemble of PDB ID 7M5T. The figure shows that most restraints are only violated in 5% or fewer of the models, suggesting the absence of systematic violations. Together with the small magnitude of the observed violations (Figures 3-4), this indicates good agreement of these experimental data with the structural models.

**Fig. 5.**
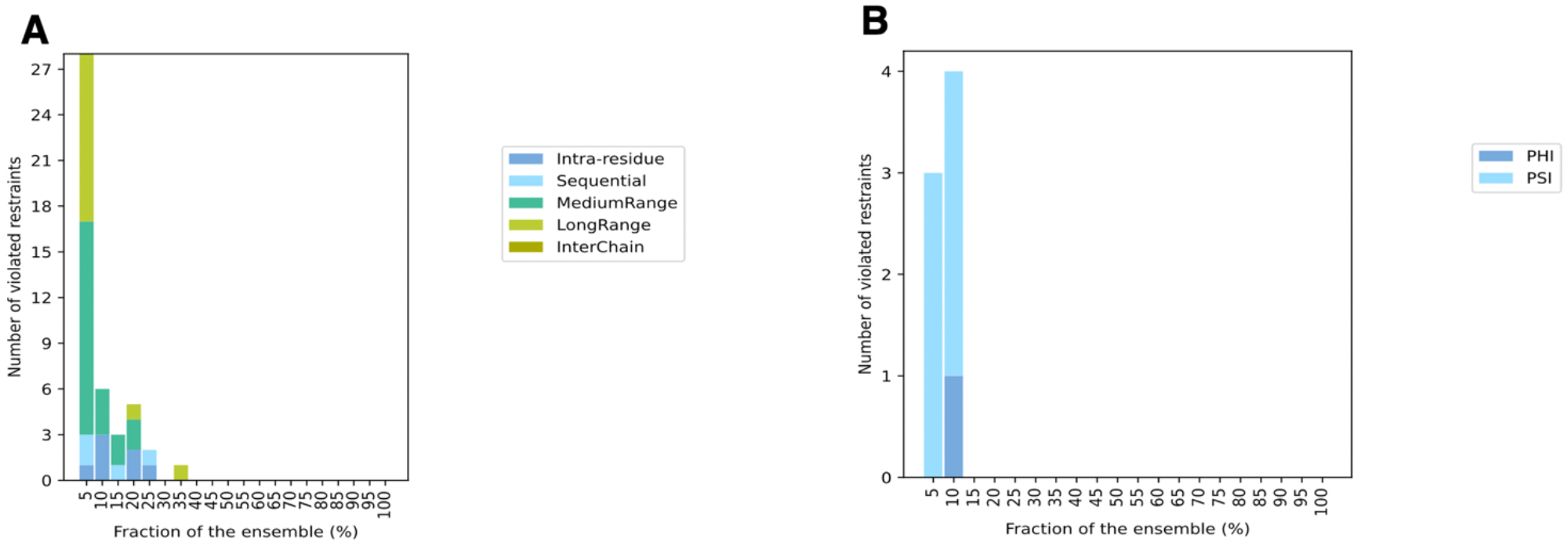
Number of (A) distance and (B) dihedral angle violations *versus* the size of the ensemble for the PDB ID 7M5T.

Histograms of each restraint’s average violation and the most violated restraints for the ensemble are given in sections 9.4 and 10.4 of the validation report for distance and dihedral-angle restraints, respectively. Similarly, sections 9.5 and 10.5 of the report provide lists of all distance and dihedral angle violations in each model in the ensemble. It also provides the histogram of the magnitude of these violations (not shown).

## DISCUSSION

In this paper, we describe a comprehensive set of new biomolecule distance- and dihedral-angle restraint validation tools for the wwPDB OneDep validation pipeline, generating both human and computer-readable reports. Although examples provided herein pertain exclusively to proteins, the same restraint validation methods can be used for distance restraint validation of other biomolecules modeled from NMR-based restraints, including nucleic acids and carbohydrates.

As most biomolecular NMR structures are determined from distance restraints (including both NOE and paramagnetic relaxation enhancement – PRE base restraints) and dihedral angle restraints, it is necessary to provide a standardized validation of models against these restraints. The current implementation of wwPDB NMR Structure Validation Software provides these tools, using upper and lower bound distance restraints, dihedral angle restraints, and chemical shift data in either NMR-STAR ver3 or NEF v1.1 data formats. In the course of this work, these features of the wwPDB NMR Validation Software have also been incorporated into the C/C++ program PDBStat ver5.23 (https://github.rpi.edu/RPIBioinformatics/PDBStat_public) (Tejero *et al*., 2013), allowing cross-validation between PDBStat and wwPDB Restraint Validation Software. This cross-validation was used to check the implementation of the rules and processes outlined in this paper. These tools are also useful in providing a standardized restraint validation protocol that can be applied across the PDB archive and used in providing standardized structure validation reports to support the publication of NMR-derived biomolecular models. In conjunction with this validation software, the CcpNmr Analysis version-3 program suite (Skinner *et al*., 2016) has also been made fully compatible with generating the required NEF input data and accepting the output of the validation pipeline for further inspection and analysis. In addition, programs for NMR structure generation, such as Xplor-NIH (Schwieters *et al*., 2006), ARIA (Rieping *et al*., 2007), and others under development, have been updated to accept both NEF and NMR-STAR formatted input data, as well as generating a consistent pair of NEF-PDBx / mmCIF formatted result files for data exchange and for OneDep deposition.

Using NEF-PDBx / mmCIF conventions and formats for defining restraints will ensure accurate and reproducible model vs. restraint data validation in the future.

Results generated using the OneDep validation pipeline, both for structure-based validation and restraint validation, are available as PDF files for human inspection and interpretation, as well as in XML and PDBx/mmCIF formats for further processing by other software programs. As a demonstration, we used CcpNmr AnalysisStructure to import and process the XML file for PDB entries 2PNG and 1PQX. We used the restraint violations to generate a per-model / per-residue metric and color-coded the structural ensembles. Fig. 6a shows the result for PDB ID 2PNG (http://doi.org/10.2210/pdb2PNG/pdb), revealing localized hot spots of substantial restraint violations that warrant further inspection. In contrast, Fig. 6b shows the result for PDB ID 1PQX (http://doi.org/10.2210/pdb1PQX/pdb), revealing very few and only incidental violations. The NEF format also provides so-called linkage information, *i.e*., between restraints and originating peaks, thus allowing for the re-examination of the originating spectral data, *e.g*., in CcpNmr AnalysisStructure.

**Fig. 6.**
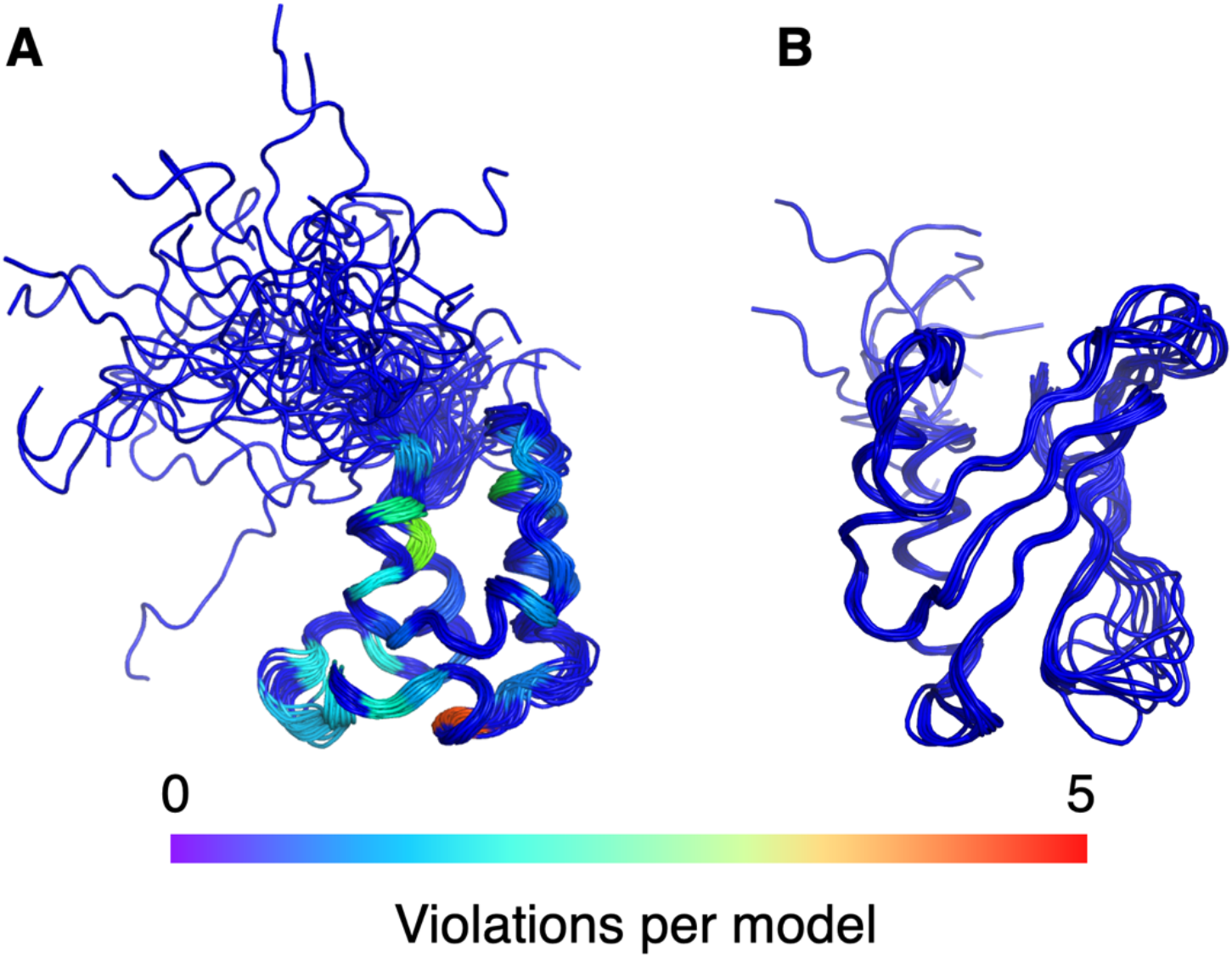
Consistent distance restraint violations mapped on the structural ensemble of (A) PDB ID 2PNG and (B) PDB ID 1PQX. Per model, per-residue restraint violations > 0.3 Å observed in > 50% of the models are mapped in blue-yellow color ramp representing 0 to 5 violated restraints per model. Violations were calculated using validation reports generated by the wwPDB validation system (validate.wwpdb.org) with coordinates in PDBx / mmCIF format and NEF formatted restraints (available at the NEF GitHub repository).

Consistency between the restraints and the model(s) is a necessary but not sufficient criterion for an accurate model. Restraints deposited by the user have often undergone a filtering and editing process during structure elucidation. Structures that exhibit no violations of the deposited restraints could still have incorrect features. Restraint violations may also result from conformational structural dynamics when modeling the biomolecule as a single conformation (*vide infra*). The validation report presented in this paper is not designed to judge the quality of the structure based on its residual restraint violations alone, but rather to show how well the model fits to the reported restraints. The Violations Report should be viewed as one of multiple tools used to validate the model, with particular value in identifying problems of consistency between the restraint list and the deposited models and/or elucidating aspects of the dynamic nature of the biomolecule.

The wwPDB is committed to improving data quality by making validation reports available to the public. Consequently, an effort to standardize existing restraints and chemical shifts into single NEF and NMR-Star formats is underway. This remediation effort will include PDB archive-wide re-generation of wwPDB validation reports with restraint validation, which will enable archival statistical assessments for outlier detection.

Although the wwPDB restraint validation software is a significant advance, other valuable model-vs-data tools are also available in the NMR spectroscopist tool chest. It is also possible to validate models against a NOE completeness metric, assessing the percentage of restraints predicted by the model that are included in the restraint list (Doreleijers *et al*., 1999). As geometrical restraints are derived from empirical NMR data, such as spectra or peak lists, various tools have also been described for the validation of models against NOESY peak list and chemical shift data (Huang *et al*., 2005; Huang *et al*., 2012; Rosato *et al*., 2013), or even directly against spectra (Thomas *et al*., 1991; Gorler and Kalbitzer, 1997; Ried *et al*., 2004).

Models can also be validated by comparing metrics of flexibility based on chemical shifts with models of flexibility derived from the structure models (Fowler *et al*., 2020; Fowler and Williamson, 2022), back calculation of chemical shifts from molecular models (Neal *et al*., 2003; Vila *et al*., 2008; Shen and Bax, 2010), or by back-calculation of residual dipolar coupling data from models (Cornilescu *et al*., 1998; Clore and Garrett, 1999; Losonczi *et al*., 1999). Each of these methods has strengths and weaknesses (Rosato *et al*., 2013). Several of these model-vs-data methods, including the NOE completeness score (Doreleijers *et al*., 1999), the RPF-DP score for assessing models against NOESY peak lists (Huang *et al*., 2005; Huang *et al*., 2012), and the RDC Q factor (Cornilescu *et al*., 1998; Clore and Garrett, 1999; Losonczi *et al*., 1999) are available as servers and are also implemented in the software package PDBStat version 5.23 (Tejero *et al*., 2013). While the use of one or more of these model-vs-data structure quality assessment methods is strongly recommended for depositors of NMR-based structural models to the wwPDB, these model-vs-data validation methods are not yet adopted by the wwPDB because there is not yet sufficient community consensus on their general applicability in the context of a global biomolecular structure archive.

Another important area of methods development involves the representation of multiple conformational states of proteins within a single PDB entry. NMR structures are generally represented by a collection of models, representing the consistency and uncertainty of atomic positions in the structural model. Each of these models is consistent with all of the available NMR data. However, in some cases, the NMR data should be more accurately interpreted in terms of multiple biomolecule conformations in dynamic equilibrium, *i.e*., the “model” should be two or more conformations present in the same sample. In the past, these multiple conformational states have been represented in various ways in PDB depositions. Future expansions of the wwPDB NMR Structure Validation software will need to account for multiple chemical shift data, multiple restraint data, and multiple atomic coordinate sets that result from multiple conformational state modeling (Ramelot *et al*., 2023). Fortunately, NEF and NMR-STAR are inherently flexible and extensible, allowing them to be implemented as a standard in these situations.

Since June 30, 2019, the wwPDB sites exclusively accept macromolecular crystallographic structures in the PDB exchange macromolecular Crystallographic Information File (PDBx / mmCIF) format (Adams *et al*., 2019). While NMR-derived structures in legacy PDB format are still accepted, the OneDep NMR Structure Validation software now also requires PDBx / mmCIF format for atomic coordinates and either NMR-STAR or NEF formats for NMR restraint files. As the requirement for providing atomic coordinates for NMR-derived structures in PDBx / mmCIF and NMR data in NMR-STAR / NEF format is anticipated in the near future, it is important that the community begin the process of adopting these formats and conventions. Both NMR-STAR and NEF also support NOESY peak list and RDC data formats, anticipating support of these data types into the validation process in the future. While validation against NOESY peak lists and RDC data are anticipated for future expansions of the wwPDB NMR Structure Validation software, additional consensus of the broader NMR community will be needed before standardizing these validation metrics.

The NMR software developer Community actively supports the development and implementation of NEF. A recent round-robin NEF testing exercise, which included the wwPDB consortium implementing the current validation pipeline, provided important insights into the practical implementation of NEF and associated challenges. A more detailed account of this exercise, including a detailed description of the NEF data format, will be presented elsewhere.

In conclusion, we presented the rationale for model-vs-data restraint validation by the wwPDB, together with a summary of validation tools for NMR distance and dihedral restraints, as implemented in the wwPDB validation pipeline and recommended by the wwPDB NMR-VTF committee (Montelione *et al*., 2013). These tools will allow for a more comprehensive, and therefore better assessment of the quality of biomolecular NMR structures and thereby benefit all users of the PDB biomolecular structure archive.

## Acknowledgments

We thank structural biologists worldwide who have contributed structures to the PDB and members of the wwPDB NMR Validation Task Force and NEF Working Group for setting data standards and validation recommendations. We also thank all the staff members of the wwPDB partners for their support and feedback and wwPDB biocurators for testing and feedback on the wwPDB validation software. RCSB PDB is funded by the National Science Foundation (DBI- 1832184), the U.S. Department of Energy (DE-SC0019749), the National Cancer Institute, the National Institute of Allergy and Infectious Diseases, and the National Institute of General Medical Sciences of the NIH under grant R01GM133198. Protein Data Bank in Europe is supported by the European Molecular Biology Laboratory-European Bioinformatics Institute and the Wellcome Trust (104948). Protein Data Bank Japan is supported by the Database Integration Coordination Program (JPMJND2205) from the department of National Bioscience Database Center (NBDC)-JST (Japan Science and Technology Agency), the Platform Project for Supporting in Drug Discovery and Life Science Research from AMED (22ama121001), and the joint usage program of Institute for Protein Research, Osaka University. BMRB is supported by the US National Institute of General Medical Sciences under grants R01GM109046 and R24GM150793. R.T., T.A.R., and G.T.M. are supported by US National Institute of General Medical Sciences grant 1R35GM141818. E.P., G.W.V., and co-workers are supported by UKRI-MRC partnership grants MR/L000555/1 and MR/P00038X/1. RCSB PDB core operations are jointly funded by the National Science Foundation (DBI-1832184, PI: S.K.B.), the US Department of Energy (DE-SC0019749, PI: S.K.B.), and the National Cancer Institute, the National Institute of Allergy and Infectious Diseases, and the National Institute of General Medical Sciences of the National Institutes of Health (R01GM133198, PI: S.K.B.). Other funding awards to RCSB PDB by the NSF and to PDBe by the UK Biotechnology and Biological Research Council are jointly supporting development of a Next Generation PDB archive (DBI- 2019297, PI: S.K.B.; BB/V004247/1, PI: S.V.) and new Mol* features (DBI-2129634, PI: S.K.B.; BB/W017970/1, PI: S.V.). CDS is supported by the Intramural Program of the National Institute of Diabetes and Digestive and Kidney Diseases. The content is solely the responsibility of the authors and does not necessarily represent the official views of the National Institutes of Health.

## Author contributions

All authors contributed to the development of the restraint validation standards outlined in this manuscript. The NEF data standards and dictionary were developed by E.A.P., G.W.V., J.W. and E.P. and is maintained by the contacts with the NEF working group developers. The corresponding PDBx / mmCIF dictionary was developed by J.W. The wwPDB restraints validation software was developed by K.B. and tested against the PDBStat software by R.T. and E.A.P. The deposition and data checking of NEF and NMR-Star data were developed by M.Y. The testing and feedback of wwPDB validation software and NEF deposition within OneDep were performed by M.C., D.H., I.P., Y.L., M.S., E.A.P., and J.T. wwPDB project management on the development of validation software and NEF data deposition was provided by J.Y.Y. Experimental data for structure validation were provided by T.A.R. The wwPDB validation software package is maintained by wwPDB partners headed by S.K.B., S.V., G.K., A.P., and J.C.H. G.T.M. and G.W.V. provided the scientific steering to the project. The article was written by K.B., G.T.M., G.W.V., and J.Y.Y., with contributions from all authors.

## Supplementary Material

wwPDB NMR Structure Validation Report for target PDB ID 7m5t

## STAR Methods

### Key resources table

Not applicable

### Resource availability

Not applicable

### Lead Contact

Further information and requests for resources should be directed to and will be fulfilled by the Lead Contact, Kumaran Baskaran.

### Materials availability

This study did not generate new unique reagents.

## Data and code availability

<u>wwPDB</u> validation tools are publicly accessible. The wwPDB anonymous validation server is provided at https://validate.wwpdb.org and the wwPDB validation API is accessible at http://www.wwpdb.org/validation/onedep-validation-web-service-interface. wwPDB validation report for each PDB ID is provided for users to download at PDB archive, https://ftp.wwpdb.org/pub/pdb/validation_reports/. These validation reports are also accessible via PDB DOI, e.g., <u>10.2210/pdb7M5T/pdb which links to its DOI landing page</u> at wwPDB website https://www.wwpdb.org/pdb?id=pdb_00007M5T.

The NEF standard and related code are available at https://github.com/NMRExchangeFormat/NEF/. The corresponding PDBx/mmCIF dictionary is accessible at https://mmcif.wwpdb.org/dictionaries/mmcif_nef.dic/Index/.

PDBStat is available under an open-source license at https://github.rpi.edu/RPIBioinformatics CcpNmr AnalysisStructure is available from https://www.ccpn.ac.uk under an open-source license for non-commercial usage.

### Experimental model and subject details

There was no model used.

### Method Details

The NMR exchange format (NEF) (Gutmanas *et al*., 2015) https://github.com/NMRExchangeFormat/NEF/) presents a community-supported standard for the interchange of NMR data between different software programs. The format is based upon the STAR syntax (Hall, 1991) and defines so-called saveframes, *i.e*., self-contained blocks of data, for sequence, chemical shifts, resonance peaks in NMR spectra, dihedral-, distance-, and RDC-restraints, as well as relevant metadata and a linkage table connecting restraints and peaks. Importantly, the format is inherently extendable through so-called namespace-specific tags, both with respect to the data contained in each saveframe or as complete additional saveframes.

The NEF defines a nomenclature convention for the twenty common protein amino acids and the eight RNA/DNA oligonucleotides in their common appearance in NMR spectra (*i.e*., appropriately protonated at pH 7.0). This nomenclature follows the IUPAC convention, with extensions to accommodate NMR-specific situations that follow from degenerate resonances and stereo-specificity, *e.g*., for methylene protons and VAL, LEU methyl groups. Key aspects of these extensions are the existence of a wild-card indicator (“%”, *e.g*., as in HB%) and indicators for non-degenerate, but non stereo-specifically assigned resonances (“x” and “y”, as in HBx and HBy). Together with the presence of a specific atom-based tag (<u>atom_site.pdbx_atom_ambiguity)</u> in the structural PDBx / mmCIF file, this allows for an unambiguous and exact mapping of the NMR restraint onto the molecular structure. A detailed description of the NEF will be presented elsewhere.

### Quantification and statistical analysis

No statistical analysis was performed.

