## Supplemental Info 1 for "Restraint Validation of Biomolecular Structures Determined by NMR in the Protein Data Bank"

### Full wwPDB NMR Structure Validation Report (i)

Aug 4, 2022 – 07:26 AM EDT

| | | |
|---|---|---|
| PDB ID | : | 7M5T |
| Title | : | Solution NMR structure of de novo designed protein 0515 |
| Authors | : | Ramelot, T.A.; Hao, J.; Baker, D.; Montelione, G.T. |
| Deposited on | : | 2021-03-24 |

This is a Full wwPDB NMR Structure Validation Report for a publicly released PDB entry.

We welcome your comments at **
A user guide is available at
https://www.wwpdb.org/validation/2017/NMRValidationReportHelp
with specific help available everywhere you see the (i) symbol.

The types of validation reports are described at http://www.wwpdb.org/validation/2017/FAQs#types.

The following versions of software and data (see references (i)) were used in the production of this report:

| | | |
|---|---|---|
| MolProbity | : | 4.02b-467 |
| Percentile statistics | : | 20191225.v01 (using entries in the PDB archive December 25th 2019) |
| RCI | : | v_1n_11_5_13_A (Berjanski et al., 2005) |
| PANAV | : | Wang et al. (2010) |
| ShiftChecker | : | 2.29 |
| BMRB Restraints Analysis | : | v1.2 |
| Ideal geometry (proteins) | : | Engh & Huber (2001) |
| Ideal geometry (DNA, RNA) | : | Parkinson et al. (1996) |
| Validation Pipeline (wwPDB-VP) | : | 2.29 |

### 1 Overall quality at a glance ⓘ

The following experimental techniques were used to determine the structure:
*SOLUTION NMR*

The overall completeness of chemical shifts assignment is 91%.

Percentile scores (ranging between 0-100) for global validation metrics of the entry are shown in the following graphic. The table shows the number of entries on which the scores are based.

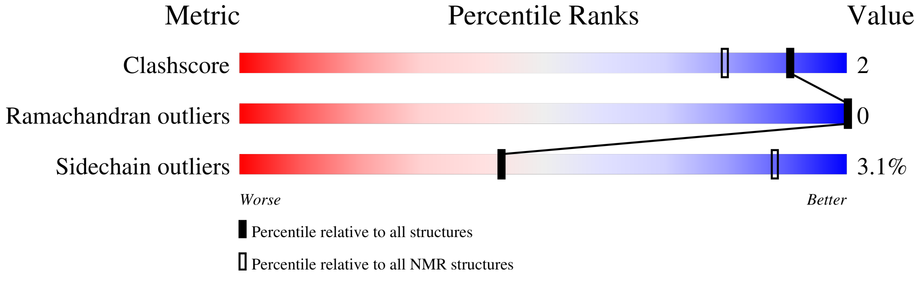


| Metric | Whole archive (#Entries) | NMR archive (#Entries) |
|---|---|---|
| Clashscore | 158937 | 12864 |
| Ramachandran outliers | 154571 | 11451 |
| Sidechain outliers | 154315 | 11428 |

The table below summarises the geometric issues observed across the polymeric chains and their fit to the experimental data. The red, orange, yellow and green segments indicate the fraction of residues that contain outliers for >=3, 2, 1 and 0 types of geometric quality criteria. A cyan segment indicates the fraction of residues that are not part of the well-defined cores, and a grey segment represents the fraction of residues that are not modelled. The numeric value for each fraction is indicated below the corresponding segment, with a dot representing fractions <=5%

| Mol | Chain | Length | Quality of chain |
|---|---|---|---|
| 1 | A | 100 | 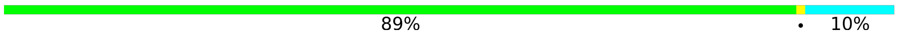  |

### 2 Ensemble composition and analysis (i)

This entry contains 20 models. Model 1 is the overall representative, medoid model (most similar to other models).

The following residues are included in the computation of the global validation metrics.

| Well-defined (core) protein residues | | | |
|---|---|---|---|
| **Well-defined core** | **Residue range (total)** | **Backbone RMSD (Å)** | **Medoid model** |
| 1 | A:3-A:70, A:79-A:100 (90) | 0.37 | 1 |

Ill-defined regions of proteins are excluded from the global statistics.

Ligands and non-protein polymers are included in the analysis.

The models can be grouped into 4 clusters. No single-model clusters were found.

| Cluster number | Models |
|---|---|
| 1 | 1, 2, 7, 11, 14, 17, 18, 19 |
| 2 | 4, 5, 8, 9, 12, 16, 20 |
| 3 | 3, 6, 13 |
| 4 | 10, 15 |

### 3 Entry composition (i)

There is only 1 type of molecule in this entry. The entry contains 1595 atoms, of which 786 are hydrogens and 0 are deuteriums.

- Molecule 1 is a protein called De novo designed protein 0515.

| Mol | Chain | Residues | Atoms | | | | | | Trace |
|---|---|---|---|---|---|---|---|---|---|
| | | | Total | C | H | N | O | S | |
| 1 | A | 100 | 1595 | 508 | 786 | 142 | 157 | 2 | 0 |

### 4 Residue-property plots (i)

#### 4.1 Average score per residue in the NMR ensemble

These plots are provided for all protein, RNA, DNA and oligosaccharide chains in the entry. The first graphic is the same as shown in the summary in section 1 of this report. The second graphic shows the sequence where residues are colour-coded according to the number of geometric quality criteria for which they contain at least one outlier: green = 0, yellow = 1, orange = 2 and red = 3 or more. Stretches of 2 or more consecutive residues without any outliers are shown as green connectors. Residues which are classified as ill-defined in the NMR ensemble, are shown in cyan with an underline colour-coded according to the previous scheme. Residues which were present in the experimental sample, but not modelled in the final structure are shown in grey.

- Molecule 1: De novo designed protein 0515

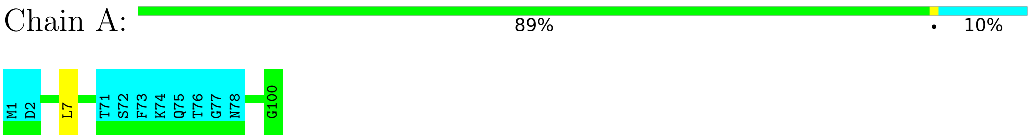


#### 4.2 Scores per residue for each member of the ensemble

Colouring as in section 4.1 above.

##### 4.2.1 Score per residue for model 1 (medoid)

- Molecule 1: De novo designed protein 0515

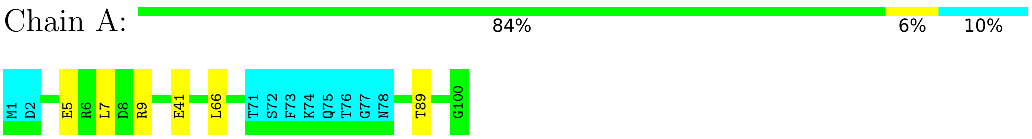


##### 4.2.2 Score per residue for model 2

- Molecule 1: De novo designed protein 0515

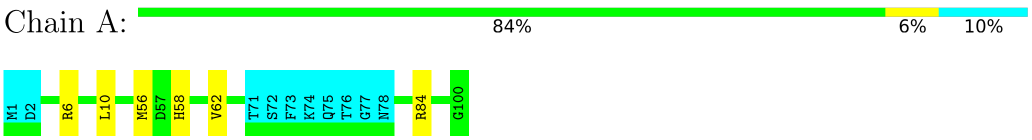

##### 4.2.3 Score per residue for model 3

- Molecule 1: De novo designed protein 0515

Chain A:

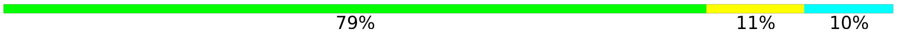


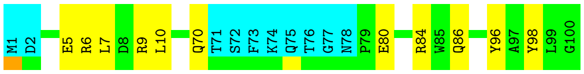


##### 4.2.4 Score per residue for model 4

- Molecule 1: De novo designed protein 0515

Chain A:

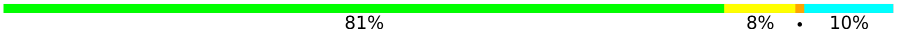


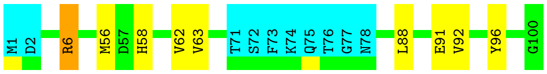


##### 4.2.5 Score per residue for model 5

- Molecule 1: De novo designed protein 0515

Chain A:

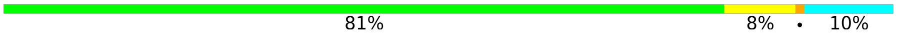


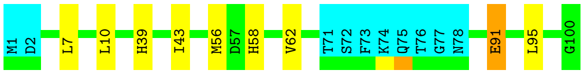


##### 4.2.6 Score per residue for model 6

- Molecule 1: De novo designed protein 0515

Chain A:

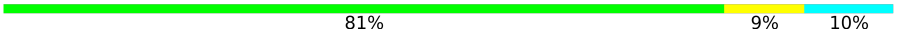


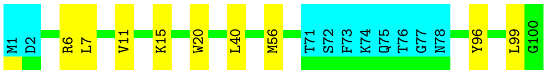


##### 4.2.7 Score per residue for model 7

- Molecule 1: De novo designed protein 0515

Chain A:

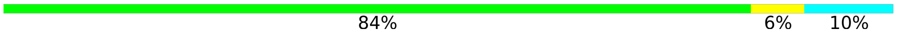


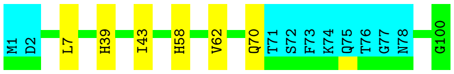

##### 4.2.8 Score per residue for model 8

- Molecule 1: De novo designed protein 0515

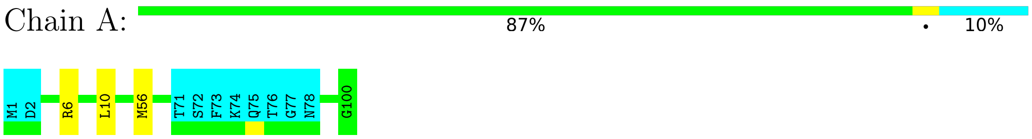


##### 4.2.9 Score per residue for model 9

- Molecule 1: De novo designed protein 0515

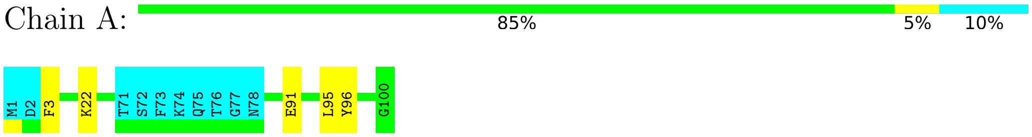


##### 4.2.10 Score per residue for model 10

- Molecule 1: De novo designed protein 0515

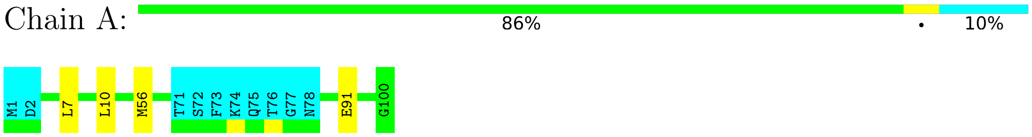


##### 4.2.11 Score per residue for model 11

- Molecule 1: De novo designed protein 0515

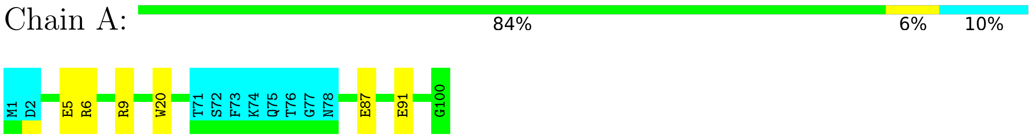


##### 4.2.12 Score per residue for model 12

- Molecule 1: De novo designed protein 0515

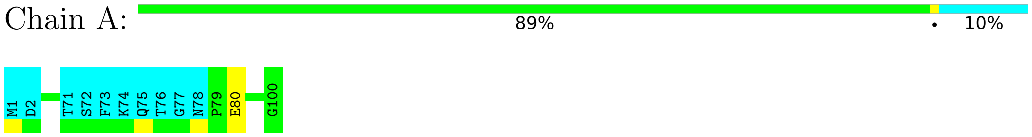

##### 4.2.13 Score per residue for model 13

- Molecule 1: De novo designed protein 0515

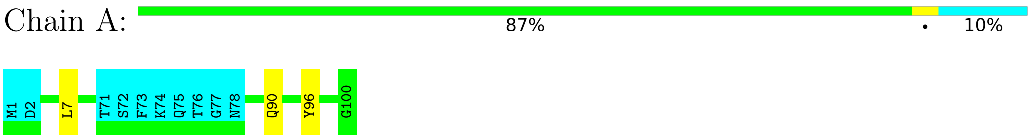


##### 4.2.14 Score per residue for model 14

- Molecule 1: De novo designed protein 0515

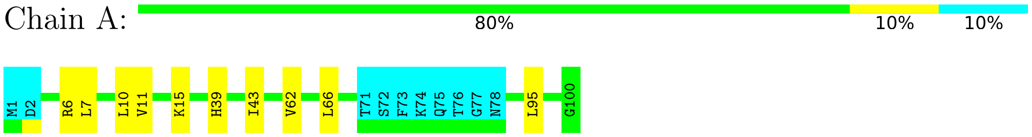


##### 4.2.15 Score per residue for model 15

- Molecule 1: De novo designed protein 0515

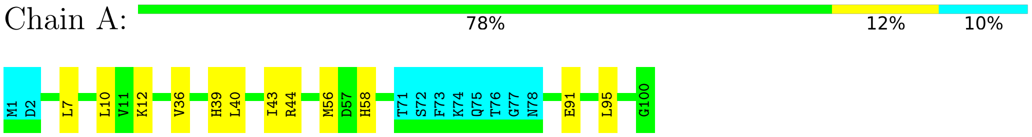


##### 4.2.16 Score per residue for model 16

- Molecule 1: De novo designed protein 0515

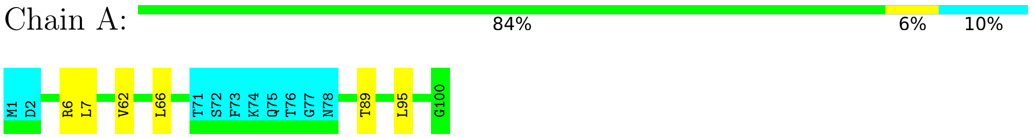


##### 4.2.17 Score per residue for model 17

- Molecule 1: De novo designed protein 0515

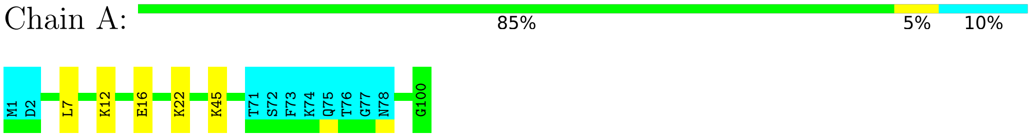

##### 4.2.18 Score per residue for model 18

• Molecule 1: De novo designed protein 0515

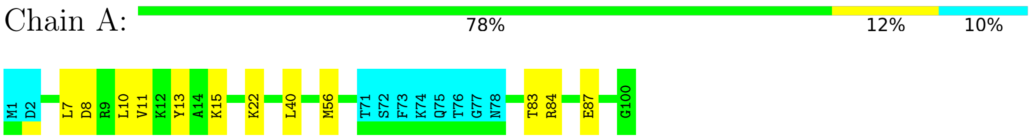


##### 4.2.19 Score per residue for model 19

• Molecule 1: De novo designed protein 0515

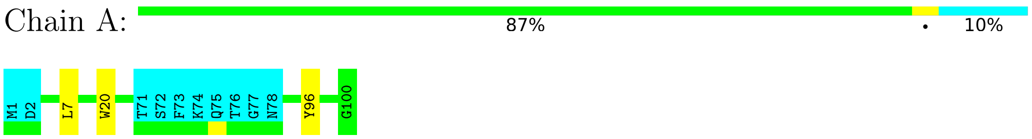


##### 4.2.20 Score per residue for model 20

• Molecule 1: De novo designed protein 0515

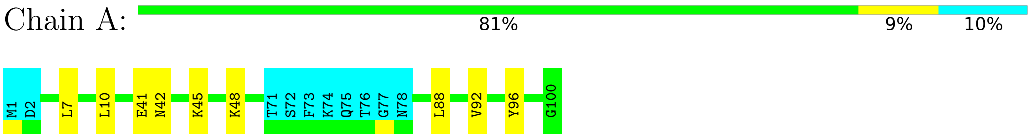

### 5 Refinement protocol and experimental data overview (i)

The models were refined using the following method: *DGSA-distance geometry simulated annealing*.

Of the 100 calculated structures, 20 were deposited, based on the following criterion: *target function*.

The following table shows the software used for structure solution, optimisation and refinement.

| Software name | Classification | Version |
|---|---|---|
| CNS | refinement | 1.3 |
| CYANA | structure calculation | 3.98.13 |
| TALOS-N | structure calculation | |

The following table shows chemical shift validation statistics as aggregates over all chemical shift files. Detailed validation can be found in section 7 of this report.

| | |
|---|---|
| Chemical shift file(s) | working_cs.cif |
| Number of chemical shift lists | 1 |
| Total number of shifts | 1243 |
| Number of shifts mapped to atoms | 1243 |
| Number of unparsed shifts | 0 |
| Number of shifts with mapping errors | 0 |
| Number of shifts with mapping warnings | 0 |
| Assignment completeness (well-defined parts) | 91% |

### 6 Model quality (i)

#### 6.1 Standard geometry (i)

There are no covalent bond-length or bond-angle outliers.

There are no bond-length outliers.

There are no bond-angle outliers.

There are no chirality outliers.

There are no planarity outliers.

#### 6.2 Too-close contacts (i)

In the following table, the Non-H and H(model) columns list the number of non-hydrogen atoms and hydrogen atoms in each chain respectively. The H(added) column lists the number of hydrogen atoms added and optimized by MolProbity. The Clashes column lists the number of clashes averaged over the ensemble.

| Mol | Chain | Non-H | H(model) | H(added) | Clashes |
|---|---|---|---|---|---|
| 1 | A | 732 | 713 | 710 | 2±2 |
| All | All | 14640 | 14260 | 14200 | 47 |

The all-atom clashscore is defined as the number of clashes found per 1000 atoms (including hydrogen atoms). The all-atom clashscore for this structure is 2.

All unique clashes are listed below, sorted by their clash magnitude.

| Atom-1 | Atom-2 | Clash(Å) | Distance(Å) | Models | |
|---|---|---|---|---|---|
| | | | | Worst | Total |
| 1:A:7:LEU:HD13 | 1:A:10:LEU:HD12 | 0.54 | 1.80 | 18 | 5 |
| 1:A:6:ARG:HA | 1:A:6:ARG:HE | 0.47 | 1.69 | 4 | 1 |
| 1:A:11:VAL:O | 1:A:15:LYS:HB2 | 0.46 | 2.10 | 14 | 3 |
| 1:A:66:LEU:HD13 | 1:A:89:THR:HA | 0.45 | 1.89 | 16 | 2 |
| 1:A:58:HIS:O | 1:A:62:VAL:HG23 | 0.44 | 2.12 | 4 | 4 |
| 1:A:56:MET:HG2 | 1:A:99:LEU:HB3 | 0.44 | 1.88 | 6 | 1 |
| 1:A:87:GLU:O | 1:A:91:GLU:HG2 | 0.44 | 2.12 | 11 | 1 |
| 1:A:91:GLU:O | 1:A:95:LEU:HD22 | 0.43 | 2.13 | 9 | 3 |
| 1:A:43:ILE:HG12 | 1:A:58:HIS:HB2 | 0.43 | 1.90 | 15 | 1 |
| 1:A:39:HIS:O | 1:A:43:ILE:HG13 | 0.43 | 2.14 | 15 | 4 |
| 1:A:6:ARG:O | 1:A:10:LEU:HG | 0.43 | 2.14 | 8 | 4 |
| 1:A:5:GLU:O | 1:A:9:ARG:HG2 | 0.43 | 2.14 | 11 | 2 |
| 1:A:80:GLU:O | 1:A:84:ARG:HG2 | 0.43 | 2.13 | 3 | 1 |
| 1:A:6:ARG:HG3 | 1:A:98:TYR:HE2 | 0.43 | 1.73 | 3 | 1 |

Continued on next page...

*Continued from previous page...*

| Atom-1 | Atom-2 | Clash(Å) | Distance(Å) | Models | |
|---|---|---|---|---|---|
| | | | | Worst | Total |
| 1:A:62:VAL:O | 1:A:66:LEU:HG | 0.42 | 2.15 | 14 | 2 |
| 1:A:63:VAL:HG21 | 1:A:96:TYR:CE1 | 0.42 | 2.50 | 4 | 1 |
| 1:A:10:LEU:HD21 | 1:A:91:GLU:HB2 | 0.42 | 1.92 | 15 | 1 |
| 1:A:88:LEU:O | 1:A:92:VAL:HG23 | 0.41 | 2.15 | 20 | 2 |
| 1:A:83:THR:O | 1:A:87:GLU:HG2 | 0.41 | 2.15 | 18 | 1 |
| 1:A:5:GLU:O | 1:A:9:ARG:HD3 | 0.41 | 2.16 | 3 | 1 |
| 1:A:36:VAL:O | 1:A:40:LEU:HG | 0.41 | 2.16 | 15 | 1 |
| 1:A:6:ARG:HB3 | 1:A:95:LEU:HD21 | 0.41 | 1.93 | 16 | 1 |
| 1:A:13:TYR:HA | 1:A:84:ARG:HD2 | 0.41 | 1.92 | 18 | 1 |
| 1:A:12:LYS:O | 1:A:16:GLU:HG2 | 0.41 | 2.16 | 17 | 1 |
| 1:A:10:LEU:HD13 | 1:A:40:LEU:HD22 | 0.40 | 1.92 | 18 | 1 |
| 1:A:7:LEU:HD11 | 1:A:40:LEU:O | 0.40 | 2.17 | 6 | 1 |

#### 6.3 Torsion angles (i)

##### 6.3.1 Protein backbone (i)

In the following table, the Percentiles column shows the percent Ramachandran outliers of the chain as a percentile score with respect to all PDB entries followed by that with respect to all NMR entries. The Analysed column shows the number of residues for which the backbone conformation was analysed and the total number of residues.

| Mol | Chain | Analysed | Favoured | Allowed | Outliers | Percentiles | |
|---|---|---|---|---|---|---|---|
| 1 | A | 89/100 (89%) | 88±1 (99±1%) | 1±1 (1±1%) | 0±0 (0±0%) | 100 | 100 |
| All | All | 1780/2000 (89%) | 1763 (99%) | 17 (1%) | 0 (0%) | 100 | 100 |

There are no Ramachandran outliers.

##### 6.3.2 Protein sidechains (i)

In the following table, the Percentiles column shows the percent sidechain outliers of the chain as a percentile score with respect to all PDB entries followed by that with respect to all NMR entries. The Analysed column shows the number of residues for which the sidechain conformation was analysed and the total number of residues.

| Mol | Chain | Analysed | Rotameric | Outliers | Percentiles | |
|---|---|---|---|---|---|---|
| 1 | A | 77/86 (90%) | 75±1 (97±1%) | 2±1 (3±1%) | 43 | 88 |
| All | All | 1540/1720 (90%) | 1493 (97%) | 47 (3%) | 43 | 88 |

All 20 unique residues with a non-rotameric sidechain are listed below. They are sorted by the frequency of occurrence in the ensemble.

| Mol | Chain | Res | Type | Models (Total) |
|---|---|---|---|---|
| 1 | A | 7 | LEU | 8 |
| 1 | A | 56 | MET | 7 |
| 1 | A | 96 | TYR | 6 |
| 1 | A | 91 | GLU | 3 |
| 1 | A | 20 | TRP | 3 |
| 1 | A | 22 | LYS | 3 |
| 1 | A | 41 | GLU | 2 |
| 1 | A | 70 | GLN | 2 |
| 1 | A | 45 | LYS | 2 |
| 1 | A | 84 | ARG | 1 |
| 1 | A | 86 | GLN | 1 |
| 1 | A | 6 | ARG | 1 |
| 1 | A | 80 | GLU | 1 |
| 1 | A | 90 | GLN | 1 |
| 1 | A | 95 | LEU | 1 |
| 1 | A | 12 | LYS | 1 |
| 1 | A | 44 | ARG | 1 |
| 1 | A | 8 | ASP | 1 |
| 1 | A | 42 | ASN | 1 |
| 1 | A | 48 | LYS | 1 |

###### 6.3.3 RNA (i)

There are no RNA molecules in this entry.

##### 6.4 Non-standard residues in protein, DNA, RNA chains (i)

There are no non-standard protein/DNA/RNA residues in this entry.

##### 6.5 Carbohydrates (i)

There are no monosaccharides in this entry.

##### 6.6 Ligand geometry (i)

There are no ligands in this entry.

#### 6.7 Other polymers (i)

There are no such molecules in this entry.

#### 6.8 Polymer linkage issues (i)

There are no chain breaks in this entry.

### 7 Chemical shift validation (i)

The completeness of assignment taking into account all chemical shift lists is 91% for the well-defined parts and 89% for the entire structure.

#### 7.1 Chemical shift list 1

File name: working_cs.cif

Chemical shift list name: *nef_chemical_shift_list_7m5t_cs.str*

##### 7.1.1 Bookkeeping (i)

The following table shows the results of parsing the chemical shift list and reports the number of nuclei with statistically unusual chemical shifts.

| | |
|---|---|
| Total number of shifts | 1243 |
| Number of shifts mapped to atoms | 1243 |
| Number of unparsed shifts | 0 |
| Number of shifts with mapping errors | 0 |
| Number of shifts with mapping warnings | 0 |
| Number of shift outliers (ShiftChecker) | 0 |

##### 7.1.2 Chemical shift referencing (i)

The following table shows the suggested chemical shift referencing corrections.

| Nucleus | # values | Correction ± precision, *ppm* | Suggested action |
|---|---|---|---|
| $^{13}C_{\alpha}$ | 98 | -0.62 ± 0.08 | Should be applied |
| $^{13}C_{\beta}$ | 93 | 0.24 ± 0.05 | None needed (< 0.5 ppm) |
| $^{13}C'$ | 94 | -0.46 ± 0.09 | None needed (< 0.5 ppm) |
| $^{15}N$ | 92 | 0.13 ± 0.11 | None needed (< 0.5 ppm) |

##### 7.1.3 Completeness of resonance assignments (i)

The following table shows the completeness of the chemical shift assignments for the well-defined regions of the structure. The overall completeness is 91%, i.e. 1043 atoms were assigned a chemical shift out of a possible 1143. 15 out of 15 assigned methyl groups (LEU and VAL) were assigned stereospecifically.

| | Total | $^{1}$H | $^{13}$C | $^{15}$N |
|---|---|---|---|---|
| Backbone | 442/444 (100%) | 177/177 (100%) | 178/180 (99%) | 87/87 (100%) |
| Sidechain | 511/592 (86%) | 313/346 (90%) | 191/215 (89%) | 7/31 (23%) |

*Continued on next page...*

*Continued from previous page...*

| | Total | $^{1}$H | $^{13}$C | $^{15}$N |
|---|---|---|---|---|
| Aromatic | 90/107 (84%) | 47/55 (85%) | 41/44 (93%) | 2/8 (25%) |
| Overall | 1043/1143 (91%) | 537/578 (93%) | 410/439 (93%) | 96/126 (76%) |

The following table shows the completeness of the chemical shift assignments for the full structure. The overall completeness is 89%, i.e. 1122 atoms were assigned a chemical shift out of a possible 1258. 15 out of 15 assigned methyl groups (LEU and VAL) were assigned stereospecifically.

| | Total | $^{1}$H | $^{13}$C | $^{15}$N |
|---|---|---|---|---|
| Backbone | 474/494 (96%) | 190/197 (96%) | 192/200 (96%) | 92/97 (95%) |
| Sidechain | 549/648 (85%) | 337/379 (89%) | 204/235 (87%) | 8/34 (24%) |
| Aromatic | 99/116 (85%) | 52/60 (87%) | 45/48 (94%) | 2/8 (25%) |
| Overall | 1122/1258 (89%) | 579/636 (91%) | 441/483 (91%) | 102/139 (73%) |

##### 7.1.4 Statistically unusual chemical shifts (i)

There are no statistically unusual chemical shifts.

##### 7.1.5 Random Coil Index (RCI) plots (i)

The image below reports *random coil index* values for the protein chains in the structure. The height of each bar gives a probability of a given residue to be disordered, as predicted from the available chemical shifts and the amino acid sequence. A value above 0.2 is an indication of significant predicted disorder. The colour of the bar shows whether the residue is in the well-defined core (black) or in the ill-defined residue ranges (cyan), as described in section 2 on ensemble composition.

Random coil index (RCI) for chain A:

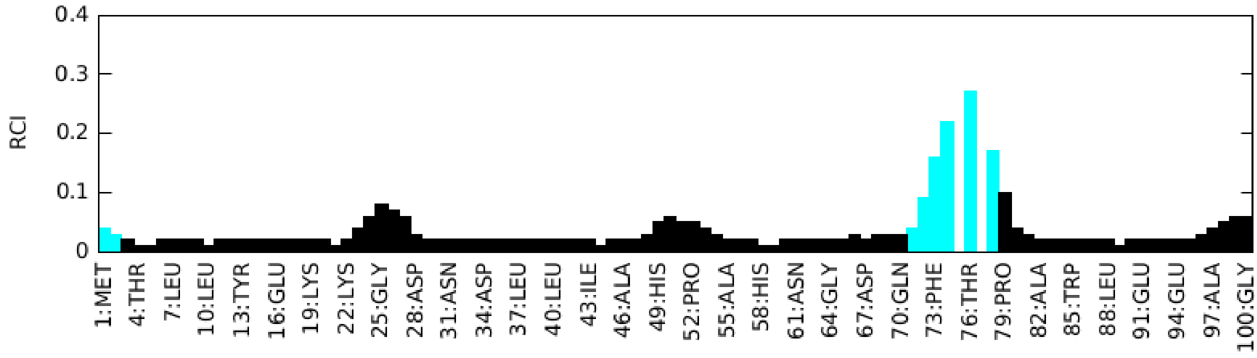

### 8 NMR restraints analysis (i)

#### 8.1 Conformationally restricting restraints (i)

The following table provides the summary of experimentally observed NMR restraints in different categories. Restraints are classified into different categories based on the sequence separation of the atoms involved.

| Description | Value |
|---|---|
| Total distance restraints | 2189 |
| Intra-residue (\|i-j\|=0) | 471 |
| Sequential (\|i-j\|=1) | 505 |
| Medium range (\|i-j\|>1 and \|i-j\|<5) | 675 |
| Long range (\|i-j\|≥5) | 398 |
| Inter-chain | 0 |
| Hydrogen bond restraints | 140 |
| Disulfide bond restraints | 0 |
| Total dihedral-angle restraints | 175 |
| Number of unmapped restraints | 0 |
| Number of restraints per residue | 23.6 |
| Number of long range restraints per residue[1] | 4.0 |

[1]Long range hydrogen bonds and disulfide bonds are counted as long range restraints while calculating the number of long range restraints per residue

#### 8.2 Residual restraint violations (i)

This section provides the overview of the restraint violations analysis. The violations are binned as small, medium and large violations based on its absolute value. Average number of violations per model is calculated by dividing the total number of violations in each bin by the size of the ensemble.

##### 8.2.1 Average number of distance violations per model (i)

Distance violations less than 0.1 Å are not included in the calculation.

| Bins (Å) | Average number of violations per model | Max (Å) |
|---|---|---|
| 0.1-0.2 (Small) | 3.6 | 0.2 |
| 0.2-0.5 (Medium) | 0.9 | 0.32 |
| >0.5 (Large) | None | None |

##### 8.2.2 Average number of dihedral-angle violations per model (i)

Dihedral-angle violations less than 1° are not included in the calculation.

| Bins (°) | Average number of violations per model | Max (°) |
|---|---|---|
| 1.0-10.0 (Small) | 0.6 | 3.5 |
| 10.0-20.0 (Medium) | None | None |
| >20.0 (Large) | None | None |

### 9 Distance violation analysis (i)

#### 9.1 Summary of distance violations (i)

The following table shows the summary of distance violations in different restraint categories based on the sequence separation of the atoms involved. Each category is further sub-divided into three sub-categories based on the atoms involved. Violations less than 0.1 Å are not included in the statistics.

| Restraints type | Count | %[1] | Violated[3] | | | Consistently Violated[4] | | |
|---|---|---|---|---|---|---|---|---|
| | | | Count | %[2] | %[1] | Count | %[2] | %[1] |
| Intra-residue (\|i-j\|=0) | 471 | 21.5 | 7 | 1.5 | 0.3 | 0 | 0.0 | 0.0 |
| Backbone-Backbone | 3 | 0.1 | 0 | 0.0 | 0.0 | 0 | 0.0 | 0.0 |
| Backbone-Sidechain | 383 | 17.5 | 6 | 1.6 | 0.3 | 0 | 0.0 | 0.0 |
| Sidechain-Sidechain | 85 | 3.9 | 1 | 1.2 | 0.0 | 0 | 0.0 | 0.0 |
| Sequential (\|i-j\|=1) | 505 | 23.1 | 4 | 0.8 | 0.2 | 0 | 0.0 | 0.0 |
| Backbone-Backbone | 97 | 4.4 | 0 | 0.0 | 0.0 | 0 | 0.0 | 0.0 |
| Backbone-Sidechain | 353 | 16.1 | 2 | 0.6 | 0.1 | 0 | 0.0 | 0.0 |
| Sidechain-Sidechain | 55 | 2.5 | 2 | 3.6 | 0.1 | 0 | 0.0 | 0.0 |
| Medium range (\|i-j\|>1 & \|i-j\|<5) | 675 | 30.8 | 21 | 3.1 | 1.0 | 0 | 0.0 | 0.0 |
| Backbone-Backbone | 231 | 10.6 | 0 | 0.0 | 0.0 | 0 | 0.0 | 0.0 |
| Backbone-Sidechain | 332 | 15.2 | 10 | 3.0 | 0.5 | 0 | 0.0 | 0.0 |
| Sidechain-Sidechain | 112 | 5.1 | 11 | 9.8 | 0.5 | 0 | 0.0 | 0.0 |
| Long range (\|i-j\|≥5) | 398 | 18.2 | 13 | 3.3 | 0.6 | 0 | 0.0 | 0.0 |
| Backbone-Backbone | 4 | 0.2 | 0 | 0.0 | 0.0 | 0 | 0.0 | 0.0 |
| Backbone-Sidechain | 161 | 7.4 | 4 | 2.5 | 0.2 | 0 | 0.0 | 0.0 |
| Sidechain-Sidechain | 233 | 10.6 | 9 | 3.9 | 0.4 | 0 | 0.0 | 0.0 |
| Inter-chain | 0 | 0.0 | 0 | 0.0 | 0.0 | 0 | 0.0 | 0.0 |
| Backbone-Backbone | 0 | 0.0 | 0 | 0.0 | 0.0 | 0 | 0.0 | 0.0 |
| Backbone-Sidechain | 0 | 0.0 | 0 | 0.0 | 0.0 | 0 | 0.0 | 0.0 |
| Sidechain-Sidechain | 0 | 0.0 | 0 | 0.0 | 0.0 | 0 | 0.0 | 0.0 |
| Hydrogen bond | 140 | 6.4 | 1 | 0.7 | 0.0 | 0 | 0.0 | 0.0 |
| Disulfide bond | 0 | 0.0 | 0 | 0.0 | 0.0 | 0 | 0.0 | 0.0 |
| Total | 2189 | 100.0 | 46 | 2.1 | 2.1 | 0 | 0.0 | 0.0 |
| Backbone-Backbone | 475 | 21.7 | 1 | 0.2 | 0.0 | 0 | 0.0 | 0.0 |
| Backbone-Sidechain | 1229 | 56.1 | 22 | 1.8 | 1.0 | 0 | 0.0 | 0.0 |
| Sidechain-Sidechain | 485 | 22.2 | 23 | 4.7 | 1.1 | 0 | 0.0 | 0.0 |

[1] percentage calculated with respect to the total number of distance restraints, [2] percentage calculated with respect to the number of restraints in a particular restraint category, [3] violated in at least one model, [4] violated in all the models

##### 9.1.1 Bar chart : Distribution of distance restraints and violations (i)

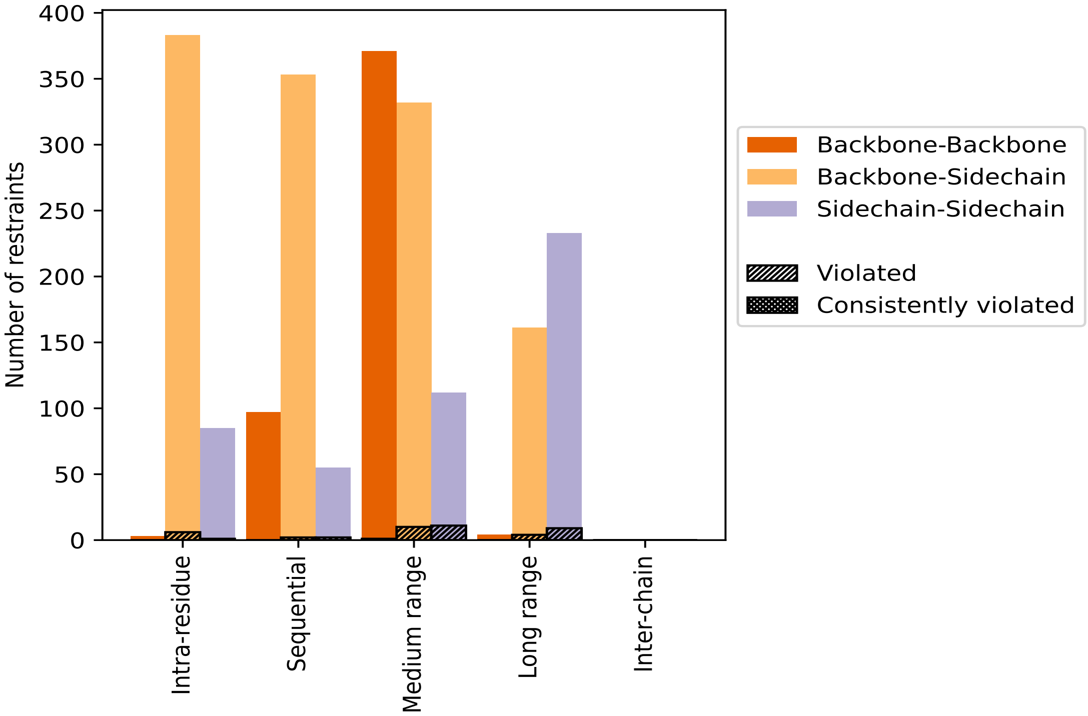


Violated and consistently violated restraints are shown using different hatch patterns in their respective categories. The hydrogen bonds and disulfied bonds are counted in their appropriate category on the x-axis

#### 9.2 Distance violation statistics for each model (i)

The following table provides the distance violation statistics for each model in the ensemble. Violations less than 0.1 Å are not included in the statistics.

| Model ID | Number of violations | | | | | | Mean (Å) | Max (Å) | SD[6] (Å) | Median (Å) |
|---|---|---|---|---|---|---|---|---|---|---|
| | IR[1] | SQ[2] | MR[3] | LR[4] | IC[5] | Total | | | | |
| 1 | 1 | 1 | 3 | 1 | 0 | 6 | 0.15 | 0.18 | 0.02 | 0.15 |
| 2 | 1 | 0 | 2 | 0 | 0 | 3 | 0.19 | 0.23 | 0.04 | 0.21 |
| 3 | 1 | 0 | 4 | 2 | 0 | 7 | 0.18 | 0.3 | 0.06 | 0.18 |
| 4 | 2 | 2 | 0 | 2 | 0 | 6 | 0.16 | 0.32 | 0.08 | 0.12 |
| 5 | 1 | 1 | 0 | 1 | 0 | 3 | 0.13 | 0.15 | 0.02 | 0.14 |
| 6 | 2 | 0 | 6 | 0 | 0 | 8 | 0.17 | 0.24 | 0.05 | 0.17 |
| 7 | 1 | 0 | 1 | 1 | 0 | 3 | 0.17 | 0.24 | 0.05 | 0.15 |
| 8 | 1 | 1 | 3 | 2 | 0 | 7 | 0.12 | 0.18 | 0.02 | 0.12 |
| 9 | 1 | 0 | 2 | 1 | 0 | 4 | 0.14 | 0.17 | 0.02 | 0.14 |
| 10 | 0 | 1 | 3 | 3 | 0 | 7 | 0.18 | 0.25 | 0.05 | 0.15 |
| 11 | 1 | 1 | 2 | 1 | 0 | 5 | 0.17 | 0.23 | 0.04 | 0.19 |

Continued on next page...

*Continued from previous page...*

| Model ID | Number of violations | | | | | | Mean (Å) | Max (Å) | SD[6] (Å) | Median (Å) |
|---|---|---|---|---|---|---|---|---|---|---|
| | IR[1] | SQ[2] | MR[3] | LR[4] | IC[5] | Total | | | | |
| 12 | 1 | 1 | 0 | 1 | 0 | 3 | 0.16 | 0.22 | 0.04 | 0.14 |
| 13 | 0 | 0 | 1 | 1 | 0 | 2 | 0.16 | 0.18 | 0.02 | 0.16 |
| 14 | 0 | 0 | 1 | 1 | 0 | 2 | 0.15 | 0.15 | 0.0 | 0.15 |
| 15 | 2 | 1 | 0 | 1 | 0 | 4 | 0.18 | 0.24 | 0.04 | 0.17 |
| 16 | 2 | 0 | 2 | 1 | 0 | 5 | 0.15 | 0.24 | 0.04 | 0.14 |
| 17 | 0 | 0 | 3 | 1 | 0 | 4 | 0.14 | 0.15 | 0.01 | 0.14 |
| 18 | 2 | 1 | 2 | 1 | 0 | 6 | 0.16 | 0.25 | 0.05 | 0.14 |
| 19 | 1 | 0 | 3 | 0 | 0 | 4 | 0.17 | 0.21 | 0.03 | 0.17 |
| 20 | 0 | 0 | 0 | 1 | 0 | 1 | 0.14 | 0.14 | 0.0 | 0.14 |

[1]Intra-residue restraints, [2]Sequential restraints, [3]Medium range restraints, [4]Long range restraints, [5]Inter-chain restraints, [6]Standard deviation

##### 9.2.1 Bar graph : Distance Violation statistics for each model (i)

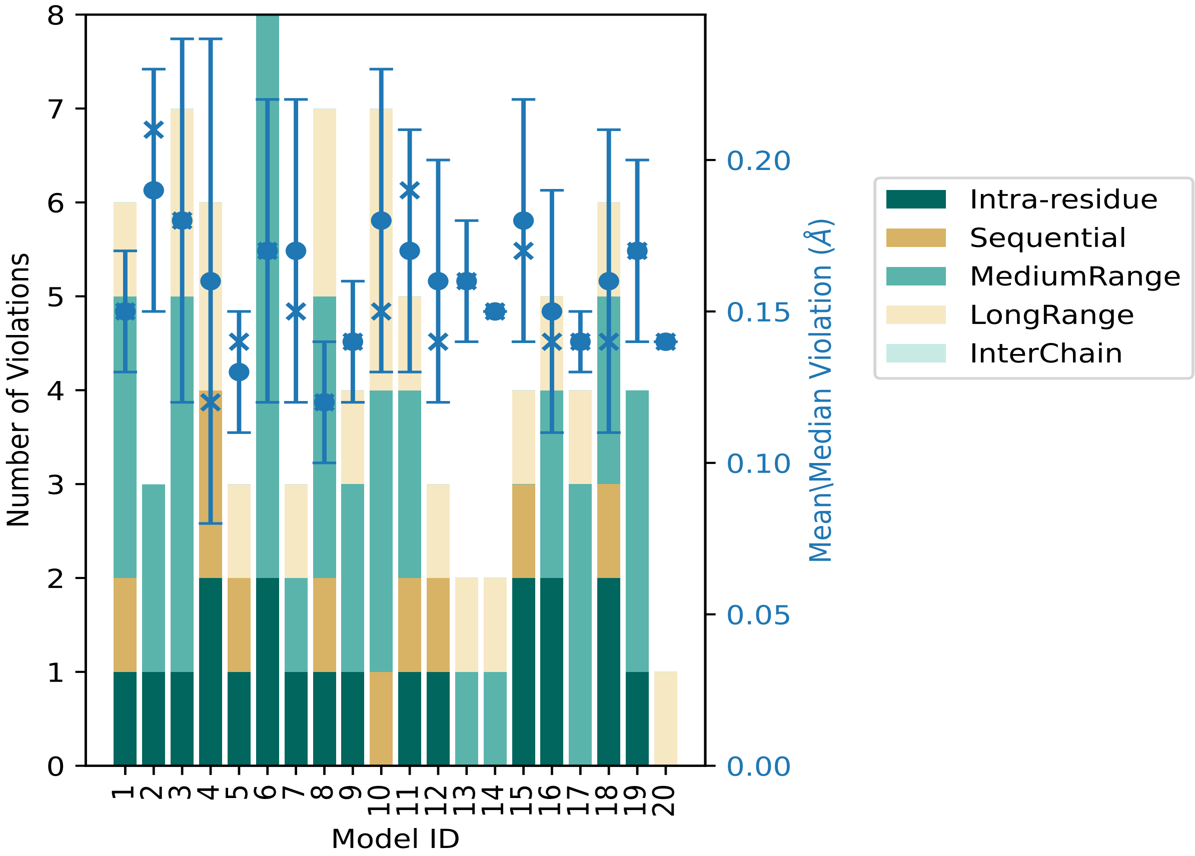


The mean(dot),median(x) and the standard deviation are shown in blue with respect to the y axis on the right

#### 9.3 Distance violation statistics for the ensemble (i)

Violation analysis may find that some restraints are violated in few models and some are violated in most of models. The following table provides this information as number of violated restraints for a given fraction of the ensemble. In total, 2004(IR:464, SQ:501, MR:654, LR:385, IC:0) restraints are not violated in the ensemble.

| Number of violated restraints | | | | | | Fraction of the ensemble | |
|---|---|---|---|---|---|---|---|
| IR[1] | SQ[2] | MR[3] | LR[4] | IC[5] | Total | Count[6] | % |
| 1 | 2 | 14 | 11 | 0 | 28 | 1 | 5.0 |
| 3 | 0 | 3 | 0 | 0 | 6 | 2 | 10.0 |
| 0 | 1 | 2 | 0 | 0 | 3 | 3 | 15.0 |
| 2 | 0 | 2 | 1 | 0 | 5 | 4 | 20.0 |
| 1 | 1 | 0 | 0 | 0 | 2 | 5 | 25.0 |
| 0 | 0 | 0 | 0 | 0 | 0 | 6 | 30.0 |
| 0 | 0 | 0 | 1 | 0 | 1 | 7 | 35.0 |
| 0 | 0 | 0 | 0 | 0 | 0 | 8 | 40.0 |
| 0 | 0 | 0 | 0 | 0 | 0 | 9 | 45.0 |
| 0 | 0 | 0 | 0 | 0 | 0 | 10 | 50.0 |
| 0 | 0 | 0 | 0 | 0 | 0 | 11 | 55.0 |
| 0 | 0 | 0 | 0 | 0 | 0 | 12 | 60.0 |
| 0 | 0 | 0 | 0 | 0 | 0 | 13 | 65.0 |
| 0 | 0 | 0 | 0 | 0 | 0 | 14 | 70.0 |
| 0 | 0 | 0 | 0 | 0 | 0 | 15 | 75.0 |
| 0 | 0 | 0 | 0 | 0 | 0 | 16 | 80.0 |
| 0 | 0 | 0 | 0 | 0 | 0 | 17 | 85.0 |
| 0 | 0 | 0 | 0 | 0 | 0 | 18 | 90.0 |
| 0 | 0 | 0 | 0 | 0 | 0 | 19 | 95.0 |
| 0 | 0 | 0 | 0 | 0 | 0 | 20 | 100.0 |

[1]Intra-residue restraints, [2]Sequential restraints, [3]Medium range restraints, [4]Long range restraints, [5]Inter-chain restraints, [6] Number of models with violations

##### 9.3.1 Bar graph : Distance violation statistics for the ensemble (i)

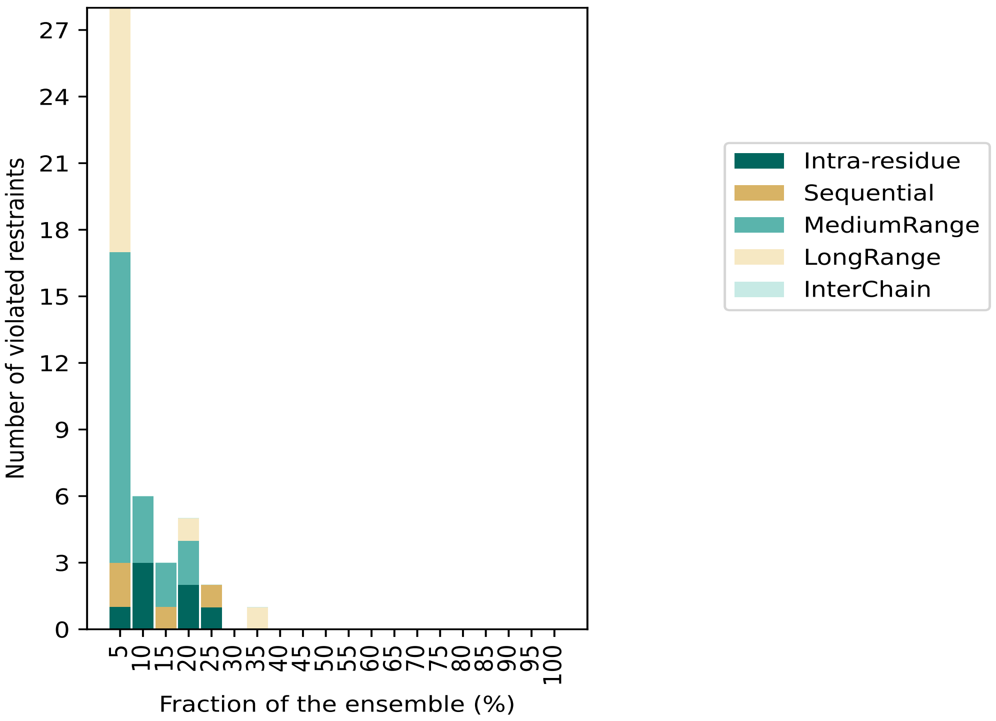


#### 9.4 Most violated distance restraints in the ensemble (i)

##### 9.4.1 Histogram : Distribution of mean distance violations (i)

The following histogram shows the distribution of the average value of the violation. The average is calculated for each restraint that is violated in more than one model over all the violated models in the ensemble

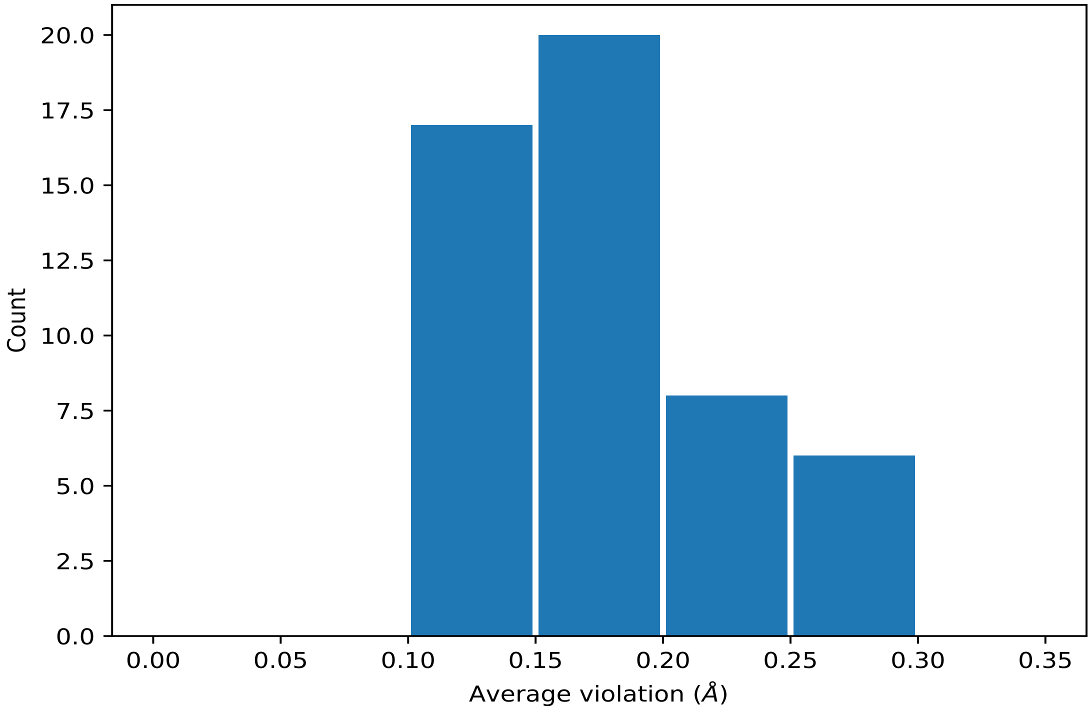


##### 9.4.2 Table: Most violated distance restraints (i)

The following table provides the mean and the standard deviation of the violation for each restraint sorted by number of violated models and the mean value. The Key (restraint list ID, restraint ID) is the unique identifier for a given restraint. Rows with same key represent combinatorial or ambiguous restraints and are counted as a single restraint.

| Key | Atom-1 | Atom-2 | Models[1] | Mean (Å) | SD[1] (Å) | Median (Å) |
|---|---|---|---|---|---|---|
| (1,325) | 1:A:8:ASP:H | 1:A:44:ARG:HG2 | 7 | 0.16 | 0.04 | 0.17 |
| (1,325) | 1:A:8:ASP:H | 1:A:44:ARG:HG3 | 7 | 0.16 | 0.04 | 0.17 |
| (1,1380) | 1:A:55:ALA:HB1 | 1:A:56:MET:HG3 | 5 | 0.15 | 0.04 | 0.14 |
| (1,1380) | 1:A:55:ALA:HB2 | 1:A:56:MET:HG3 | 5 | 0.15 | 0.04 | 0.14 |
| (1,1380) | 1:A:55:ALA:HB3 | 1:A:56:MET:HG3 | 5 | 0.15 | 0.04 | 0.14 |
| (1,1404) | 1:A:56:MET:HB2 | 1:A:56:MET:HE1 | 5 | 0.13 | 0.01 | 0.13 |
| (1,1404) | 1:A:56:MET:HB2 | 1:A:56:MET:HE2 | 5 | 0.13 | 0.01 | 0.13 |
| (1,1404) | 1:A:56:MET:HB2 | 1:A:56:MET:HE3 | 5 | 0.13 | 0.01 | 0.13 |
| (1,1404) | 1:A:56:MET:HB3 | 1:A:56:MET:HE1 | 5 | 0.13 | 0.01 | 0.13 |
| (1,1404) | 1:A:56:MET:HB3 | 1:A:56:MET:HE2 | 5 | 0.13 | 0.01 | 0.13 |
| (1,1404) | 1:A:56:MET:HB3 | 1:A:56:MET:HE3 | 5 | 0.13 | 0.01 | 0.13 |
| (1,1100) | 1:A:41:GLU:H | 1:A:41:GLU:HG2 | 4 | 0.24 | 0.0 | 0.24 |
| (1,1100) | 1:A:41:GLU:H | 1:A:41:GLU:HG3 | 4 | 0.24 | 0.0 | 0.24 |
| (1,932) | 1:A:34:ASP:HA | 1:A:37:LEU:HD21 | 4 | 0.16 | 0.04 | 0.16 |
| (1,932) | 1:A:34:ASP:HA | 1:A:37:LEU:HD22 | 4 | 0.16 | 0.04 | 0.16 |
| (1,932) | 1:A:34:ASP:HA | 1:A:37:LEU:HD23 | 4 | 0.16 | 0.04 | 0.16 |

*Continued on next page...*

*Continued from previous page...*

| Key | Atom-1 | Atom-2 | Models[1] | Mean (Å) | SD[1] (Å) | Median (Å) |
|---|---|---|---|---|---|---|
| (1,1022) | 1:A:37:LEU:H | 1:A:37:LEU:HG | 4 | 0.16 | 0.02 | 0.15 |
| (1,2041) | 1:A:92:VAL:HA | 1:A:95:LEU:HD11 | 4 | 0.15 | 0.01 | 0.15 |
| (1,2041) | 1:A:92:VAL:HA | 1:A:95:LEU:HD12 | 4 | 0.15 | 0.01 | 0.15 |
| (1,2041) | 1:A:92:VAL:HA | 1:A:95:LEU:HD13 | 4 | 0.15 | 0.01 | 0.15 |
| (1,611) | 1:A:17:ILE:HD11 | 1:A:85:TRP:HE3 | 4 | 0.14 | 0.01 | 0.14 |
| (1,611) | 1:A:17:ILE:HD12 | 1:A:85:TRP:HE3 | 4 | 0.14 | 0.01 | 0.14 |
| (1,611) | 1:A:17:ILE:HD13 | 1:A:85:TRP:HE3 | 4 | 0.14 | 0.01 | 0.14 |
| (1,137) | 1:A:95:LEU:O | 1:A:99:LEU:H | 4 | 0.13 | 0.01 | 0.12 |
| (1,340) | 1:A:9:ARG:HD2 | 1:A:12:LYS:HB2 | 3 | 0.18 | 0.04 | 0.19 |
| (1,340) | 1:A:9:ARG:HD2 | 1:A:12:LYS:HB3 | 3 | 0.18 | 0.04 | 0.19 |
| (1,340) | 1:A:9:ARG:HD3 | 1:A:12:LYS:HB2 | 3 | 0.18 | 0.04 | 0.19 |
| (1,340) | 1:A:9:ARG:HD3 | 1:A:12:LYS:HB3 | 3 | 0.18 | 0.04 | 0.19 |
| (1,915) | 1:A:33:VAL:HG11 | 1:A:37:LEU:HG | 3 | 0.14 | 0.02 | 0.13 |
| (1,915) | 1:A:33:VAL:HG12 | 1:A:37:LEU:HG | 3 | 0.14 | 0.02 | 0.13 |
| (1,915) | 1:A:33:VAL:HG13 | 1:A:37:LEU:HG | 3 | 0.14 | 0.02 | 0.13 |
| (1,1887) | 1:A:83:THR:H | 1:A:84:ARG:HG2 | 3 | 0.12 | 0.0 | 0.12 |
| (1,1887) | 1:A:83:THR:H | 1:A:84:ARG:HG3 | 3 | 0.12 | 0.0 | 0.12 |
| (1,2108) | 1:A:95:LEU:HD11 | 1:A:98:TYR:HE1 | 2 | 0.27 | 0.03 | 0.27 |
| (1,2108) | 1:A:95:LEU:HD11 | 1:A:98:TYR:HE2 | 2 | 0.27 | 0.03 | 0.27 |
| (1,2108) | 1:A:95:LEU:HD12 | 1:A:98:TYR:HE1 | 2 | 0.27 | 0.03 | 0.27 |
| (1,2108) | 1:A:95:LEU:HD12 | 1:A:98:TYR:HE2 | 2 | 0.27 | 0.03 | 0.27 |
| (1,2108) | 1:A:95:LEU:HD13 | 1:A:98:TYR:HE1 | 2 | 0.27 | 0.03 | 0.27 |
| (1,2108) | 1:A:95:LEU:HD13 | 1:A:98:TYR:HE2 | 2 | 0.27 | 0.03 | 0.27 |
| (1,563) | 1:A:15:LYS:H | 1:A:15:LYS:HD2 | 2 | 0.22 | 0.01 | 0.22 |
| (1,563) | 1:A:15:LYS:H | 1:A:15:LYS:HD3 | 2 | 0.22 | 0.01 | 0.22 |
| (1,1096) | 1:A:41:GLU:HG2 | 1:A:45:LYS:HE2 | 2 | 0.2 | 0.03 | 0.2 |
| (1,1096) | 1:A:41:GLU:HG2 | 1:A:45:LYS:HE3 | 2 | 0.2 | 0.03 | 0.2 |
| (1,1096) | 1:A:41:GLU:HG3 | 1:A:45:LYS:HE2 | 2 | 0.2 | 0.03 | 0.2 |
| (1,1096) | 1:A:41:GLU:HG3 | 1:A:45:LYS:HE3 | 2 | 0.2 | 0.03 | 0.2 |
| (1,823) | 1:A:27:PRO:HG2 | 1:A:29:PHE:H | 2 | 0.18 | 0.03 | 0.18 |
| (1,823) | 1:A:27:PRO:HG3 | 1:A:29:PHE:H | 2 | 0.18 | 0.03 | 0.18 |
| (1,1369) | 1:A:54:ARG:H | 1:A:54:ARG:HG2 | 2 | 0.15 | 0.04 | 0.15 |
| (1,1369) | 1:A:54:ARG:H | 1:A:54:ARG:HG3 | 2 | 0.15 | 0.04 | 0.15 |
| (1,771) | 1:A:22:LYS:H | 1:A:22:LYS:HG2 | 2 | 0.12 | 0.0 | 0.12 |
| (1,771) | 1:A:22:LYS:H | 1:A:22:LYS:HG3 | 2 | 0.12 | 0.0 | 0.12 |

[1]Number of violated models, [2]Standard deviation

#### 9.5 All violated distance restraints (i)

##### 9.5.1 Histogram : Distribution of distance violations (i)

The following histogram shows the distribution of the absolute value of the violation for all violated restraints in the ensemble.

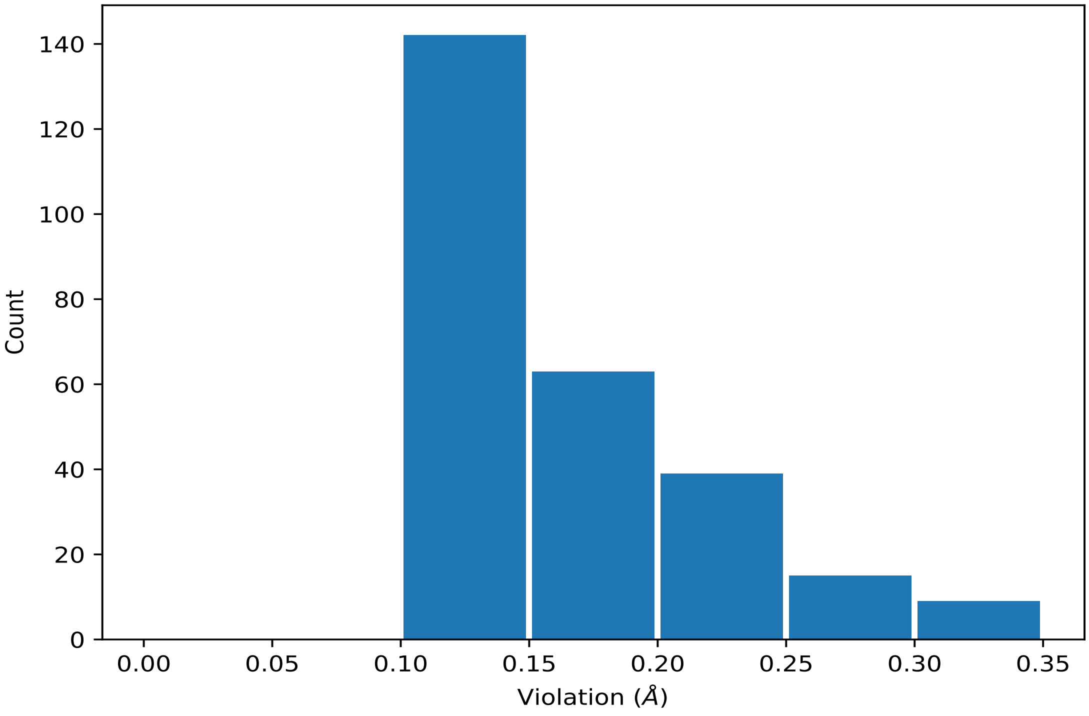


##### 9.5.2 Table : All distance violations (i)

The following table lists the absolute value of the violation for each restraint in the ensemble sorted by its value. The Key (restraint list ID, restraint ID) is the unique identifier for a given restraint. Rows with same key represent combinatorial or ambiguous restraints and are counted as a single restraint.

| Key | Atom-1 | Atom-2 | Model ID | Violation (Å) |
|---|---|---|---|---|
| (1,1603) | 1:A:63:VAL:HG21 | 1:A:96:TYR:HB3 | 4 | 0.32 |
| (1,1603) | 1:A:63:VAL:HG22 | 1:A:96:TYR:HB3 | 4 | 0.32 |
| (1,1603) | 1:A:63:VAL:HG23 | 1:A:96:TYR:HB3 | 4 | 0.32 |
| (1,2108) | 1:A:95:LEU:HD11 | 1:A:98:TYR:HE1 | 3 | 0.3 |
| (1,2108) | 1:A:95:LEU:HD11 | 1:A:98:TYR:HE2 | 3 | 0.3 |
| (1,2108) | 1:A:95:LEU:HD12 | 1:A:98:TYR:HE1 | 3 | 0.3 |
| (1,2108) | 1:A:95:LEU:HD12 | 1:A:98:TYR:HE2 | 3 | 0.3 |
| (1,2108) | 1:A:95:LEU:HD13 | 1:A:98:TYR:HE1 | 3 | 0.3 |
| (1,2108) | 1:A:95:LEU:HD13 | 1:A:98:TYR:HE2 | 3 | 0.3 |
| (1,422) | 1:A:11:VAL:HG21 | 1:A:15:LYS:HE2 | 10 | 0.25 |

*Continued on next page...*

*Continued from previous page...*

| Key | Atom-1 | Atom-2 | Model ID | Violation (Å) |
|---|---|---|---|---|
| (1,422) | 1:A:11:VAL:HG21 | 1:A:15:LYS:HE3 | 10 | 0.25 |
| (1,422) | 1:A:11:VAL:HG22 | 1:A:15:LYS:HE2 | 10 | 0.25 |
| (1,422) | 1:A:11:VAL:HG22 | 1:A:15:LYS:HE3 | 10 | 0.25 |
| (1,422) | 1:A:11:VAL:HG23 | 1:A:15:LYS:HE2 | 10 | 0.25 |
| (1,422) | 1:A:11:VAL:HG23 | 1:A:15:LYS:HE3 | 10 | 0.25 |
| (1,416) | 1:A:11:VAL:HG11 | 1:A:15:LYS:HE2 | 10 | 0.25 |
| (1,416) | 1:A:11:VAL:HG12 | 1:A:15:LYS:HE2 | 10 | 0.25 |
| (1,416) | 1:A:11:VAL:HG13 | 1:A:15:LYS:HE2 | 10 | 0.25 |
| (1,414) | 1:A:11:VAL:HG11 | 1:A:15:LYS:HD2 | 18 | 0.25 |
| (1,414) | 1:A:11:VAL:HG11 | 1:A:15:LYS:HD3 | 18 | 0.25 |
| (1,414) | 1:A:11:VAL:HG12 | 1:A:15:LYS:HD2 | 18 | 0.25 |
| (1,414) | 1:A:11:VAL:HG12 | 1:A:15:LYS:HD3 | 18 | 0.25 |
| (1,414) | 1:A:11:VAL:HG13 | 1:A:15:LYS:HD2 | 18 | 0.25 |
| (1,414) | 1:A:11:VAL:HG13 | 1:A:15:LYS:HD3 | 18 | 0.25 |
| (1,2108) | 1:A:95:LEU:HD11 | 1:A:98:TYR:HE1 | 6 | 0.24 |
| (1,2108) | 1:A:95:LEU:HD11 | 1:A:98:TYR:HE2 | 6 | 0.24 |
| (1,2108) | 1:A:95:LEU:HD12 | 1:A:98:TYR:HE1 | 6 | 0.24 |
| (1,2108) | 1:A:95:LEU:HD12 | 1:A:98:TYR:HE2 | 6 | 0.24 |
| (1,2108) | 1:A:95:LEU:HD13 | 1:A:98:TYR:HE1 | 6 | 0.24 |
| (1,2108) | 1:A:95:LEU:HD13 | 1:A:98:TYR:HE2 | 6 | 0.24 |
| (1,1100) | 1:A:41:GLU:H | 1:A:41:GLU:HG2 | 6 | 0.24 |
| (1,1100) | 1:A:41:GLU:H | 1:A:41:GLU:HG3 | 6 | 0.24 |
| (1,1100) | 1:A:41:GLU:H | 1:A:41:GLU:HG2 | 7 | 0.24 |
| (1,1100) | 1:A:41:GLU:H | 1:A:41:GLU:HG3 | 7 | 0.24 |
| (1,1100) | 1:A:41:GLU:H | 1:A:41:GLU:HG2 | 15 | 0.24 |
| (1,1100) | 1:A:41:GLU:H | 1:A:41:GLU:HG3 | 15 | 0.24 |
| (1,1100) | 1:A:41:GLU:H | 1:A:41:GLU:HG2 | 16 | 0.24 |
| (1,1100) | 1:A:41:GLU:H | 1:A:41:GLU:HG3 | 16 | 0.24 |
| (1,563) | 1:A:15:LYS:H | 1:A:15:LYS:HD2 | 3 | 0.23 |
| (1,563) | 1:A:15:LYS:H | 1:A:15:LYS:HD3 | 3 | 0.23 |
| (1,340) | 1:A:9:ARG:HD2 | 1:A:12:LYS:HB2 | 2 | 0.23 |
| (1,340) | 1:A:9:ARG:HD2 | 1:A:12:LYS:HB3 | 2 | 0.23 |
| (1,340) | 1:A:9:ARG:HD3 | 1:A:12:LYS:HB2 | 2 | 0.23 |
| (1,340) | 1:A:9:ARG:HD3 | 1:A:12:LYS:HB3 | 2 | 0.23 |
| (1,1380) | 1:A:55:ALA:HB1 | 1:A:56:MET:HG3 | 18 | 0.23 |
| (1,1380) | 1:A:55:ALA:HB2 | 1:A:56:MET:HG3 | 18 | 0.23 |
| (1,1380) | 1:A:55:ALA:HB3 | 1:A:56:MET:HG3 | 18 | 0.23 |
| (1,1096) | 1:A:41:GLU:HG2 | 1:A:45:LYS:HE2 | 11 | 0.23 |
| (1,1096) | 1:A:41:GLU:HG2 | 1:A:45:LYS:HE3 | 11 | 0.23 |
| (1,1096) | 1:A:41:GLU:HG3 | 1:A:45:LYS:HE2 | 11 | 0.23 |
| (1,1096) | 1:A:41:GLU:HG3 | 1:A:45:LYS:HE3 | 11 | 0.23 |
| (1,563) | 1:A:15:LYS:H | 1:A:15:LYS:HD2 | 12 | 0.22 |

*Continued on next page...*

*Continued from previous page...*

| Key | Atom-1 | Atom-2 | Model ID | Violation (Å) |
|---|---|---|---|---|
| (1,563) | 1:A:15:LYS:H | 1:A:15:LYS:HD3 | 12 | 0.22 |
| (1,932) | 1:A:34:ASP:HA | 1:A:37:LEU:HD21 | 19 | 0.21 |
| (1,932) | 1:A:34:ASP:HA | 1:A:37:LEU:HD22 | 19 | 0.21 |
| (1,932) | 1:A:34:ASP:HA | 1:A:37:LEU:HD23 | 19 | 0.21 |
| (1,823) | 1:A:27:PRO:HG2 | 1:A:29:PHE:H | 2 | 0.21 |
| (1,823) | 1:A:27:PRO:HG3 | 1:A:29:PHE:H | 2 | 0.21 |
| (1,325) | 1:A:8:ASP:H | 1:A:44:ARG:HG2 | 10 | 0.21 |
| (1,325) | 1:A:8:ASP:H | 1:A:44:ARG:HG3 | 10 | 0.21 |
| (1,163) | 1:A:3:PHE:HA | 1:A:95:LEU:HD11 | 11 | 0.2 |
| (1,163) | 1:A:3:PHE:HA | 1:A:95:LEU:HD12 | 11 | 0.2 |
| (1,163) | 1:A:3:PHE:HA | 1:A:95:LEU:HD13 | 11 | 0.2 |
| (1,340) | 1:A:9:ARG:HD2 | 1:A:12:LYS:HB2 | 6 | 0.19 |
| (1,340) | 1:A:9:ARG:HD2 | 1:A:12:LYS:HB3 | 6 | 0.19 |
| (1,340) | 1:A:9:ARG:HD3 | 1:A:12:LYS:HB2 | 6 | 0.19 |
| (1,340) | 1:A:9:ARG:HD3 | 1:A:12:LYS:HB3 | 6 | 0.19 |
| (1,325) | 1:A:8:ASP:H | 1:A:44:ARG:HG2 | 3 | 0.19 |
| (1,325) | 1:A:8:ASP:H | 1:A:44:ARG:HG3 | 3 | 0.19 |
| (1,1961) | 1:A:87:GLU:H | 1:A:89:THR:HB | 6 | 0.19 |
| (1,1369) | 1:A:54:ARG:H | 1:A:54:ARG:HG2 | 11 | 0.19 |
| (1,1369) | 1:A:54:ARG:H | 1:A:54:ARG:HG3 | 11 | 0.19 |
| (1,1022) | 1:A:37:LEU:H | 1:A:37:LEU:HG | 19 | 0.19 |
| (1,932) | 1:A:34:ASP:HA | 1:A:37:LEU:HD21 | 1 | 0.18 |
| (1,932) | 1:A:34:ASP:HA | 1:A:37:LEU:HD22 | 1 | 0.18 |
| (1,932) | 1:A:34:ASP:HA | 1:A:37:LEU:HD23 | 1 | 0.18 |
| (1,325) | 1:A:8:ASP:H | 1:A:44:ARG:HG2 | 15 | 0.18 |
| (1,325) | 1:A:8:ASP:H | 1:A:44:ARG:HG3 | 15 | 0.18 |
| (1,1880) | 1:A:83:THR:HG21 | 1:A:86:GLN:HE22 | 3 | 0.18 |
| (1,1880) | 1:A:83:THR:HG22 | 1:A:86:GLN:HE22 | 3 | 0.18 |
| (1,1880) | 1:A:83:THR:HG23 | 1:A:86:GLN:HE22 | 3 | 0.18 |
| (1,1236) | 1:A:46:ALA:H | 1:A:58:HIS:HD2 | 8 | 0.18 |
| (1,1096) | 1:A:41:GLU:HG2 | 1:A:45:LYS:HE2 | 13 | 0.18 |
| (1,1096) | 1:A:41:GLU:HG2 | 1:A:45:LYS:HE3 | 13 | 0.18 |
| (1,1096) | 1:A:41:GLU:HG3 | 1:A:45:LYS:HE2 | 13 | 0.18 |
| (1,1096) | 1:A:41:GLU:HG3 | 1:A:45:LYS:HE3 | 13 | 0.18 |
| (1,915) | 1:A:33:VAL:HG11 | 1:A:37:LEU:HG | 1 | 0.17 |
| (1,915) | 1:A:33:VAL:HG12 | 1:A:37:LEU:HG | 1 | 0.17 |
| (1,915) | 1:A:33:VAL:HG13 | 1:A:37:LEU:HG | 1 | 0.17 |
| (1,325) | 1:A:8:ASP:H | 1:A:44:ARG:HG2 | 9 | 0.17 |
| (1,325) | 1:A:8:ASP:H | 1:A:44:ARG:HG3 | 9 | 0.17 |
| (1,1380) | 1:A:55:ALA:HB1 | 1:A:56:MET:HG3 | 15 | 0.16 |
| (1,1380) | 1:A:55:ALA:HB2 | 1:A:56:MET:HG3 | 15 | 0.16 |
| (1,1380) | 1:A:55:ALA:HB3 | 1:A:56:MET:HG3 | 15 | 0.16 |

*Continued on next page...*

*Continued from previous page...*

| Key | Atom-1 | Atom-2 | Model ID | Violation (Å) |
|---|---|---|---|---|
| (1,823) | 1:A:27:PRO:HG2 | 1:A:29:PHE:H | 17 | 0.15 |
| (1,823) | 1:A:27:PRO:HG3 | 1:A:29:PHE:H | 17 | 0.15 |
| (1,611) | 1:A:17:ILE:HD11 | 1:A:85:TRP:HE3 | 14 | 0.15 |
| (1,611) | 1:A:17:ILE:HD12 | 1:A:85:TRP:HE3 | 14 | 0.15 |
| (1,611) | 1:A:17:ILE:HD13 | 1:A:85:TRP:HE3 | 14 | 0.15 |
| (1,2084) | 1:A:94:GLU:HA | 1:A:98:TYR:HE1 | 9 | 0.15 |
| (1,2084) | 1:A:94:GLU:HA | 1:A:98:TYR:HE2 | 9 | 0.15 |
| (1,2041) | 1:A:92:VAL:HA | 1:A:95:LEU:HD11 | 16 | 0.15 |
| (1,2041) | 1:A:92:VAL:HA | 1:A:95:LEU:HD12 | 16 | 0.15 |
| (1,2041) | 1:A:92:VAL:HA | 1:A:95:LEU:HD13 | 16 | 0.15 |
| (1,2041) | 1:A:92:VAL:HA | 1:A:95:LEU:HD11 | 17 | 0.15 |
| (1,2041) | 1:A:92:VAL:HA | 1:A:95:LEU:HD12 | 17 | 0.15 |
| (1,2041) | 1:A:92:VAL:HA | 1:A:95:LEU:HD13 | 17 | 0.15 |
| (1,2041) | 1:A:92:VAL:HA | 1:A:95:LEU:HD11 | 19 | 0.15 |
| (1,2041) | 1:A:92:VAL:HA | 1:A:95:LEU:HD12 | 19 | 0.15 |
| (1,2041) | 1:A:92:VAL:HA | 1:A:95:LEU:HD13 | 19 | 0.15 |
| (1,1413) | 1:A:56:MET:HE1 | 1:A:96:TYR:HD1 | 7 | 0.15 |
| (1,1413) | 1:A:56:MET:HE1 | 1:A:96:TYR:HD2 | 7 | 0.15 |
| (1,1413) | 1:A:56:MET:HE2 | 1:A:96:TYR:HD1 | 7 | 0.15 |
| (1,1413) | 1:A:56:MET:HE2 | 1:A:96:TYR:HD2 | 7 | 0.15 |
| (1,1413) | 1:A:56:MET:HE3 | 1:A:96:TYR:HD1 | 7 | 0.15 |
| (1,1413) | 1:A:56:MET:HE3 | 1:A:96:TYR:HD2 | 7 | 0.15 |
| (1,1404) | 1:A:56:MET:HB2 | 1:A:56:MET:HE1 | 5 | 0.15 |
| (1,1404) | 1:A:56:MET:HB2 | 1:A:56:MET:HE2 | 5 | 0.15 |
| (1,1404) | 1:A:56:MET:HB2 | 1:A:56:MET:HE3 | 5 | 0.15 |
| (1,1404) | 1:A:56:MET:HB3 | 1:A:56:MET:HE1 | 5 | 0.15 |
| (1,1404) | 1:A:56:MET:HB3 | 1:A:56:MET:HE2 | 5 | 0.15 |
| (1,1404) | 1:A:56:MET:HB3 | 1:A:56:MET:HE3 | 5 | 0.15 |
| (1,137) | 1:A:95:LEU:O | 1:A:99:LEU:H | 14 | 0.15 |
| (1,1088) | 1:A:41:GLU:HA | 1:A:44:ARG:HB2 | 10 | 0.15 |
| (1,1022) | 1:A:37:LEU:H | 1:A:37:LEU:HG | 1 | 0.15 |
| (1,1022) | 1:A:37:LEU:H | 1:A:37:LEU:HG | 6 | 0.15 |
| (1,611) | 1:A:17:ILE:HD11 | 1:A:85:TRP:HE3 | 4 | 0.14 |
| (1,611) | 1:A:17:ILE:HD12 | 1:A:85:TRP:HE3 | 4 | 0.14 |
| (1,611) | 1:A:17:ILE:HD13 | 1:A:85:TRP:HE3 | 4 | 0.14 |
| (1,611) | 1:A:17:ILE:HD11 | 1:A:85:TRP:HE3 | 16 | 0.14 |
| (1,611) | 1:A:17:ILE:HD12 | 1:A:85:TRP:HE3 | 16 | 0.14 |
| (1,611) | 1:A:17:ILE:HD13 | 1:A:85:TRP:HE3 | 16 | 0.14 |
| (1,558) | 1:A:15:LYS:HE2 | 1:A:37:LEU:HD21 | 10 | 0.14 |
| (1,558) | 1:A:15:LYS:HE2 | 1:A:37:LEU:HD22 | 10 | 0.14 |
| (1,558) | 1:A:15:LYS:HE2 | 1:A:37:LEU:HD23 | 10 | 0.14 |
| (1,558) | 1:A:15:LYS:HE3 | 1:A:37:LEU:HD21 | 10 | 0.14 |

*Continued on next page...*

*Continued from previous page...*

| Key | Atom-1 | Atom-2 | Model ID | Violation (Å) |
|---|---|---|---|---|
| (1,558) | 1:A:15:LYS:HE3 | 1:A:37:LEU:HD22 | 10 | 0.14 |
| (1,558) | 1:A:15:LYS:HE3 | 1:A:37:LEU:HD23 | 10 | 0.14 |
| (1,557) | 1:A:15:LYS:HD2 | 1:A:37:LEU:HD21 | 12 | 0.14 |
| (1,557) | 1:A:15:LYS:HD2 | 1:A:37:LEU:HD22 | 12 | 0.14 |
| (1,557) | 1:A:15:LYS:HD2 | 1:A:37:LEU:HD23 | 12 | 0.14 |
| (1,557) | 1:A:15:LYS:HD3 | 1:A:37:LEU:HD21 | 12 | 0.14 |
| (1,557) | 1:A:15:LYS:HD3 | 1:A:37:LEU:HD22 | 12 | 0.14 |
| (1,557) | 1:A:15:LYS:HD3 | 1:A:37:LEU:HD23 | 12 | 0.14 |
| (1,325) | 1:A:8:ASP:H | 1:A:44:ARG:HG2 | 20 | 0.14 |
| (1,325) | 1:A:8:ASP:H | 1:A:44:ARG:HG3 | 20 | 0.14 |
| (1,283) | 1:A:7:LEU:HD11 | 1:A:8:ASP:HA | 4 | 0.14 |
| (1,283) | 1:A:7:LEU:HD12 | 1:A:8:ASP:HA | 4 | 0.14 |
| (1,283) | 1:A:7:LEU:HD13 | 1:A:8:ASP:HA | 4 | 0.14 |
| (1,2096) | 1:A:95:LEU:HA | 1:A:98:TYR:HD1 | 6 | 0.14 |
| (1,2096) | 1:A:95:LEU:HA | 1:A:98:TYR:HD2 | 6 | 0.14 |
| (1,1949) | 1:A:87:GLU:HA | 1:A:90:GLN:HE21 | 1 | 0.14 |
| (1,1949) | 1:A:87:GLU:HA | 1:A:90:GLN:HE22 | 1 | 0.14 |
| (1,1380) | 1:A:55:ALA:HB1 | 1:A:56:MET:HG3 | 5 | 0.14 |
| (1,1380) | 1:A:55:ALA:HB2 | 1:A:56:MET:HG3 | 5 | 0.14 |
| (1,1380) | 1:A:55:ALA:HB3 | 1:A:56:MET:HG3 | 5 | 0.14 |
| (1,1095) | 1:A:41:GLU:HG2 | 1:A:45:LYS:HD2 | 19 | 0.14 |
| (1,1095) | 1:A:41:GLU:HG2 | 1:A:45:LYS:HD3 | 19 | 0.14 |
| (1,1095) | 1:A:41:GLU:HG3 | 1:A:45:LYS:HD2 | 19 | 0.14 |
| (1,1095) | 1:A:41:GLU:HG3 | 1:A:45:LYS:HD3 | 19 | 0.14 |
| (1,1022) | 1:A:37:LEU:H | 1:A:37:LEU:HG | 18 | 0.14 |
| (1,932) | 1:A:34:ASP:HA | 1:A:37:LEU:HD21 | 3 | 0.13 |
| (1,932) | 1:A:34:ASP:HA | 1:A:37:LEU:HD22 | 3 | 0.13 |
| (1,932) | 1:A:34:ASP:HA | 1:A:37:LEU:HD23 | 3 | 0.13 |
| (1,915) | 1:A:33:VAL:HG11 | 1:A:37:LEU:HG | 18 | 0.13 |
| (1,915) | 1:A:33:VAL:HG12 | 1:A:37:LEU:HG | 18 | 0.13 |
| (1,915) | 1:A:33:VAL:HG13 | 1:A:37:LEU:HG | 18 | 0.13 |
| (1,611) | 1:A:17:ILE:HD11 | 1:A:85:TRP:HE3 | 13 | 0.13 |
| (1,611) | 1:A:17:ILE:HD12 | 1:A:85:TRP:HE3 | 13 | 0.13 |
| (1,611) | 1:A:17:ILE:HD13 | 1:A:85:TRP:HE3 | 13 | 0.13 |
| (1,457) | 1:A:12:LYS:HE2 | 1:A:13:TYR:HE1 | 1 | 0.13 |
| (1,457) | 1:A:12:LYS:HE2 | 1:A:13:TYR:HE2 | 1 | 0.13 |
| (1,457) | 1:A:12:LYS:HE3 | 1:A:13:TYR:HE1 | 1 | 0.13 |
| (1,457) | 1:A:12:LYS:HE3 | 1:A:13:TYR:HE2 | 1 | 0.13 |
| (1,379) | 1:A:10:LEU:HD21 | 1:A:13:TYR:HE1 | 11 | 0.13 |
| (1,379) | 1:A:10:LEU:HD21 | 1:A:13:TYR:HE2 | 11 | 0.13 |
| (1,379) | 1:A:10:LEU:HD22 | 1:A:13:TYR:HE1 | 11 | 0.13 |
| (1,379) | 1:A:10:LEU:HD22 | 1:A:13:TYR:HE2 | 11 | 0.13 |

*Continued on next page...*

*Continued from previous page...*

| Key | Atom-1 | Atom-2 | Model ID | Violation (Å) |
|---|---|---|---|---|
| (1,379) | 1:A:10:LEU:HD23 | 1:A:13:TYR:HE1 | 11 | 0.13 |
| (1,379) | 1:A:10:LEU:HD23 | 1:A:13:TYR:HE2 | 11 | 0.13 |
| (1,340) | 1:A:9:ARG:HD2 | 1:A:12:LYS:HB2 | 9 | 0.13 |
| (1,340) | 1:A:9:ARG:HD2 | 1:A:12:LYS:HB3 | 9 | 0.13 |
| (1,340) | 1:A:9:ARG:HD3 | 1:A:12:LYS:HB2 | 9 | 0.13 |
| (1,340) | 1:A:9:ARG:HD3 | 1:A:12:LYS:HB3 | 9 | 0.13 |
| (1,2041) | 1:A:92:VAL:HA | 1:A:95:LEU:HD11 | 7 | 0.13 |
| (1,2041) | 1:A:92:VAL:HA | 1:A:95:LEU:HD12 | 7 | 0.13 |
| (1,2041) | 1:A:92:VAL:HA | 1:A:95:LEU:HD13 | 7 | 0.13 |
| (1,1737) | 1:A:70:GLN:HG2 | 1:A:85:TRP:HE1 | 17 | 0.13 |
| (1,1737) | 1:A:70:GLN:HG3 | 1:A:85:TRP:HE1 | 17 | 0.13 |
| (1,1656) | 1:A:66:LEU:HD11 | 1:A:89:THR:HA | 1 | 0.13 |
| (1,1656) | 1:A:66:LEU:HD12 | 1:A:89:THR:HA | 1 | 0.13 |
| (1,1656) | 1:A:66:LEU:HD13 | 1:A:89:THR:HA | 1 | 0.13 |
| (1,1404) | 1:A:56:MET:HB2 | 1:A:56:MET:HE1 | 2 | 0.13 |
| (1,1404) | 1:A:56:MET:HB2 | 1:A:56:MET:HE2 | 2 | 0.13 |
| (1,1404) | 1:A:56:MET:HB2 | 1:A:56:MET:HE3 | 2 | 0.13 |
| (1,1404) | 1:A:56:MET:HB3 | 1:A:56:MET:HE1 | 2 | 0.13 |
| (1,1404) | 1:A:56:MET:HB3 | 1:A:56:MET:HE2 | 2 | 0.13 |
| (1,1404) | 1:A:56:MET:HB3 | 1:A:56:MET:HE3 | 2 | 0.13 |
| (1,1404) | 1:A:56:MET:HB2 | 1:A:56:MET:HE1 | 15 | 0.13 |
| (1,1404) | 1:A:56:MET:HB2 | 1:A:56:MET:HE2 | 15 | 0.13 |
| (1,1404) | 1:A:56:MET:HB2 | 1:A:56:MET:HE3 | 15 | 0.13 |
| (1,1404) | 1:A:56:MET:HB3 | 1:A:56:MET:HE1 | 15 | 0.13 |
| (1,1404) | 1:A:56:MET:HB3 | 1:A:56:MET:HE2 | 15 | 0.13 |
| (1,1404) | 1:A:56:MET:HB3 | 1:A:56:MET:HE3 | 15 | 0.13 |
| (1,1404) | 1:A:56:MET:HB2 | 1:A:56:MET:HE1 | 18 | 0.13 |
| (1,1404) | 1:A:56:MET:HB2 | 1:A:56:MET:HE2 | 18 | 0.13 |
| (1,1404) | 1:A:56:MET:HB2 | 1:A:56:MET:HE3 | 18 | 0.13 |
| (1,1404) | 1:A:56:MET:HB3 | 1:A:56:MET:HE1 | 18 | 0.13 |
| (1,1404) | 1:A:56:MET:HB3 | 1:A:56:MET:HE2 | 18 | 0.13 |
| (1,1404) | 1:A:56:MET:HB3 | 1:A:56:MET:HE3 | 18 | 0.13 |
| (1,1380) | 1:A:55:ALA:HB1 | 1:A:56:MET:HG3 | 10 | 0.13 |
| (1,1380) | 1:A:55:ALA:HB2 | 1:A:56:MET:HG3 | 10 | 0.13 |
| (1,1380) | 1:A:55:ALA:HB3 | 1:A:56:MET:HG3 | 10 | 0.13 |
| (1,137) | 1:A:95:LEU:O | 1:A:99:LEU:H | 3 | 0.13 |
| (1,932) | 1:A:34:ASP:HA | 1:A:37:LEU:HD21 | 8 | 0.12 |
| (1,932) | 1:A:34:ASP:HA | 1:A:37:LEU:HD22 | 8 | 0.12 |
| (1,932) | 1:A:34:ASP:HA | 1:A:37:LEU:HD23 | 8 | 0.12 |
| (1,771) | 1:A:22:LYS:H | 1:A:22:LYS:HG2 | 9 | 0.12 |
| (1,771) | 1:A:22:LYS:H | 1:A:22:LYS:HG3 | 9 | 0.12 |
| (1,742) | 1:A:21:TYR:HD1 | 1:A:33:VAL:HG21 | 8 | 0.12 |

*Continued on next page...*

*Continued from previous page...*

| Key | Atom-1 | Atom-2 | Model ID | Violation (Å) |
|---|---|---|---|---|
| (1,742) | 1:A:21:TYR:HD1 | 1:A:33:VAL:HG22 | 8 | 0.12 |
| (1,742) | 1:A:21:TYR:HD1 | 1:A:33:VAL:HG23 | 8 | 0.12 |
| (1,742) | 1:A:21:TYR:HD2 | 1:A:33:VAL:HG21 | 8 | 0.12 |
| (1,742) | 1:A:21:TYR:HD2 | 1:A:33:VAL:HG22 | 8 | 0.12 |
| (1,742) | 1:A:21:TYR:HD2 | 1:A:33:VAL:HG23 | 8 | 0.12 |
| (1,389) | 1:A:10:LEU:HD21 | 1:A:95:LEU:HG | 10 | 0.12 |
| (1,389) | 1:A:10:LEU:HD22 | 1:A:95:LEU:HG | 10 | 0.12 |
| (1,389) | 1:A:10:LEU:HD23 | 1:A:95:LEU:HG | 10 | 0.12 |
| (1,216) | 1:A:4:THR:HG21 | 1:A:8:ASP:H | 17 | 0.12 |
| (1,216) | 1:A:4:THR:HG22 | 1:A:8:ASP:H | 17 | 0.12 |
| (1,216) | 1:A:4:THR:HG23 | 1:A:8:ASP:H | 17 | 0.12 |
| (1,1902) | 1:A:84:ARG:H | 1:A:84:ARG:HD2 | 16 | 0.12 |
| (1,1887) | 1:A:83:THR:H | 1:A:84:ARG:HG2 | 8 | 0.12 |
| (1,1887) | 1:A:83:THR:H | 1:A:84:ARG:HG3 | 8 | 0.12 |
| (1,1887) | 1:A:83:THR:H | 1:A:84:ARG:HG2 | 11 | 0.12 |
| (1,1887) | 1:A:83:THR:H | 1:A:84:ARG:HG3 | 11 | 0.12 |
| (1,1887) | 1:A:83:THR:H | 1:A:84:ARG:HG2 | 12 | 0.12 |
| (1,1887) | 1:A:83:THR:H | 1:A:84:ARG:HG3 | 12 | 0.12 |
| (1,137) | 1:A:95:LEU:O | 1:A:99:LEU:H | 6 | 0.12 |
| (1,1361) | 1:A:54:ARG:HD2 | 1:A:57:ASP:H | 16 | 0.12 |
| (1,1361) | 1:A:54:ARG:HD3 | 1:A:57:ASP:H | 16 | 0.12 |
| (1,915) | 1:A:33:VAL:HG11 | 1:A:37:LEU:HG | 6 | 0.11 |
| (1,915) | 1:A:33:VAL:HG12 | 1:A:37:LEU:HG | 6 | 0.11 |
| (1,915) | 1:A:33:VAL:HG13 | 1:A:37:LEU:HG | 6 | 0.11 |
| (1,771) | 1:A:22:LYS:H | 1:A:22:LYS:HG2 | 4 | 0.11 |
| (1,771) | 1:A:22:LYS:H | 1:A:22:LYS:HG3 | 4 | 0.11 |
| (1,325) | 1:A:8:ASP:H | 1:A:44:ARG:HG2 | 5 | 0.11 |
| (1,325) | 1:A:8:ASP:H | 1:A:44:ARG:HG3 | 5 | 0.11 |
| (1,325) | 1:A:8:ASP:H | 1:A:44:ARG:HG2 | 18 | 0.11 |
| (1,325) | 1:A:8:ASP:H | 1:A:44:ARG:HG3 | 18 | 0.11 |
| (1,186) | 1:A:3:PHE:HE1 | 1:A:98:TYR:HB2 | 3 | 0.11 |
| (1,186) | 1:A:3:PHE:HE1 | 1:A:98:TYR:HB3 | 3 | 0.11 |
| (1,186) | 1:A:3:PHE:HE2 | 1:A:98:TYR:HB2 | 3 | 0.11 |
| (1,186) | 1:A:3:PHE:HE2 | 1:A:98:TYR:HB3 | 3 | 0.11 |
| (1,1404) | 1:A:56:MET:HB2 | 1:A:56:MET:HE1 | 4 | 0.11 |
| (1,1404) | 1:A:56:MET:HB2 | 1:A:56:MET:HE2 | 4 | 0.11 |
| (1,1404) | 1:A:56:MET:HB2 | 1:A:56:MET:HE3 | 4 | 0.11 |
| (1,1404) | 1:A:56:MET:HB3 | 1:A:56:MET:HE1 | 4 | 0.11 |
| (1,1404) | 1:A:56:MET:HB3 | 1:A:56:MET:HE2 | 4 | 0.11 |
| (1,1404) | 1:A:56:MET:HB3 | 1:A:56:MET:HE3 | 4 | 0.11 |
| (1,1380) | 1:A:55:ALA:HB1 | 1:A:56:MET:HG3 | 4 | 0.11 |
| (1,1380) | 1:A:55:ALA:HB2 | 1:A:56:MET:HG3 | 4 | 0.11 |

*Continued on next page...*

*Continued from previous page...*

| Key | Atom-1 | Atom-2 | Model ID | Violation (Å) |
|---|---|---|---|---|
| (1,1380) | 1:A:55:ALA:HB3 | 1:A:56:MET:HG3 | 4 | 0.11 |
| (1,137) | 1:A:95:LEU:O | 1:A:99:LEU:H | 8 | 0.11 |
| (1,1369) | 1:A:54:ARG:H | 1:A:54:ARG:HG2 | 8 | 0.11 |
| (1,1369) | 1:A:54:ARG:H | 1:A:54:ARG:HG3 | 8 | 0.11 |
| (1,1198) | 1:A:45:LYS:HE2 | 1:A:49:HIS:HE1 | 8 | 0.11 |
| (1,1198) | 1:A:45:LYS:HE3 | 1:A:49:HIS:HE1 | 8 | 0.11 |

### 10 Dihedral-angle violation analysis (i)

#### 10.1 Summary of dihedral-angle violations (i)

The following table provides the summary of dihedral-angle violations in different dihedral-angle types. Violations less than 1° are not included in the calculation.

| Angle type | Count | %[1] | Violated[3] | | | Consistently Violated[4] | | |
|---|---|---|---|---|---|---|---|---|
| | | | Count | %[2] | %[1] | Count | %[2] | %[1] |
| PSI | 89 | 50.9 | 6 | 6.7 | 3.4 | 0 | 0.0 | 0.0 |
| PHI | 86 | 49.1 | 1 | 1.2 | 0.6 | 0 | 0.0 | 0.0 |
| Total | 175 | 100.0 | 7 | 4.0 | 4.0 | 0 | 0.0 | 0.0 |

[1] percentage calculated with respect to total number of dihedral-angle restraints, [2] percentage calculated with respect to number of restraints in a particular dihedral-angle type, [3] violated in at least one model, [4] violated in all the models

##### 10.1.1 Bar chart : Distribution of dihedral-angles and violations (i)

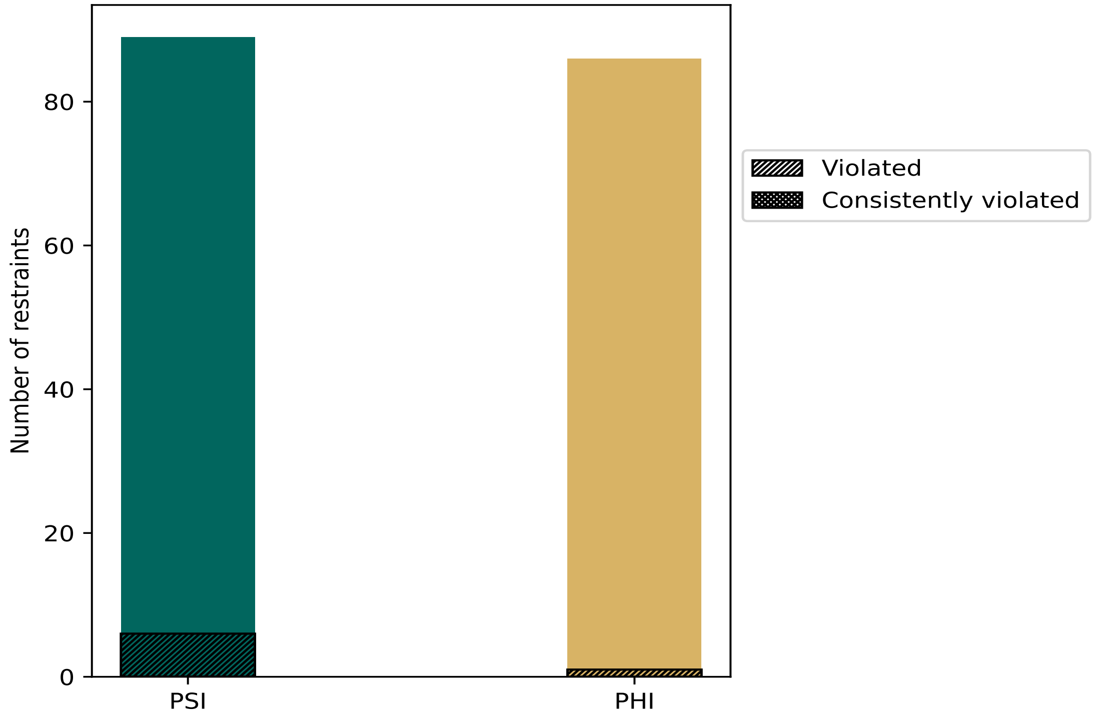


Violated and consistently violated restraints are shown using different hatch patterns in their respective categories

#### 10.2 Dihedral-angle violation statistics for each model (i)

The following table provides the dihedral-angle violation statistics for each model in the ensemble. Violations less than 1° are not included in the statistics.

| Model ID | Number of violations | | | Mean (°) | Max (°) | SD (°) | Median (°) |
|---|---|---|---|---|---|---|---|
| | PSI | PHI | Total | | | | |
| 1 | 0 | 0 | 0 | 0.0 | 0.0 | 0.0 | 0.0 |
| 2 | 1 | 0 | 1 | 1.3 | 1.3 | 0.0 | 1.3 |
| 3 | 2 | 0 | 2 | 2.25 | 2.8 | 0.55 | 2.25 |
| 4 | 0 | 0 | 0 | 0.0 | 0.0 | 0.0 | 0.0 |
| 5 | 0 | 0 | 0 | 0.0 | 0.0 | 0.0 | 0.0 |
| 6 | 2 | 0 | 2 | 2.6 | 3.5 | 0.9 | 2.6 |
| 7 | 0 | 0 | 0 | 0.0 | 0.0 | 0.0 | 0.0 |
| 8 | 1 | 0 | 1 | 1.1 | 1.1 | 0.0 | 1.1 |
| 9 | 1 | 0 | 1 | 2.3 | 2.3 | 0.0 | 2.3 |
| 10 | 0 | 1 | 1 | 1.3 | 1.3 | 0.0 | 1.3 |
| 11 | 0 | 0 | 0 | 0.0 | 0.0 | 0.0 | 0.0 |
| 12 | 0 | 0 | 0 | 0.0 | 0.0 | 0.0 | 0.0 |
| 13 | 1 | 0 | 1 | 1.1 | 1.1 | 0.0 | 1.1 |
| 14 | 0 | 1 | 1 | 1.6 | 1.6 | 0.0 | 1.6 |
| 15 | 0 | 0 | 0 | 0.0 | 0.0 | 0.0 | 0.0 |
| 16 | 0 | 0 | 0 | 0.0 | 0.0 | 0.0 | 0.0 |
| 17 | 0 | 0 | 0 | 0.0 | 0.0 | 0.0 | 0.0 |
| 18 | 0 | 0 | 0 | 0.0 | 0.0 | 0.0 | 0.0 |
| 19 | 1 | 0 | 1 | 1.4 | 1.4 | 0.0 | 1.4 |
| 20 | 0 | 0 | 0 | 0.0 | 0.0 | 0.0 | 0.0 |

##### 10.2.1 Bar graph : Dihedral violation statistics for each model (i)

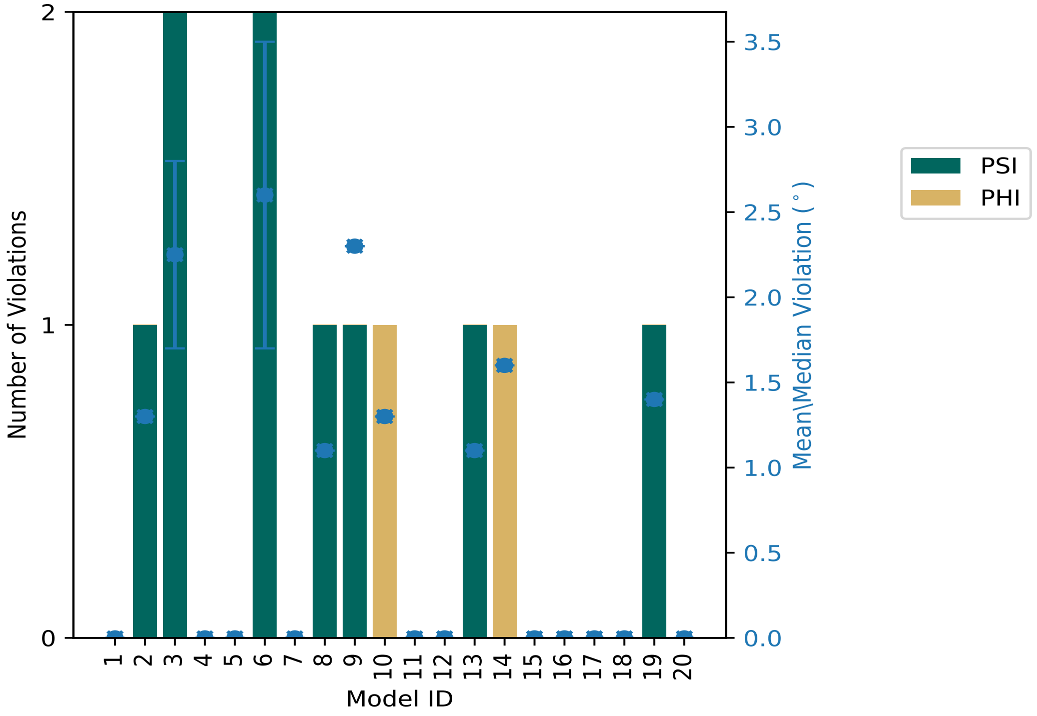


The mean(dot),median(x) and the standard deviation are shown in blue with respect to the y axis on the right

#### 10.3 Dihedral-angle violation statistics for the ensemble (i)

Violation analysis may find that some restraints are violated in very few models and some are violated in most of models. The following table provides this information as number of violated restraints for a given fraction of ensemble.

| Number of violated restraints | | | Fraction of the ensemble | |
|---|---|---|---|---|
| PSI | PHI | Total | Count[1] | % |
| 3 | 0 | 3 | 1 | 5.0 |
| 3 | 1 | 4 | 2 | 10.0 |
| 0 | 0 | 0 | 3 | 15.0 |
| 0 | 0 | 0 | 4 | 20.0 |
| 0 | 0 | 0 | 5 | 25.0 |
| 0 | 0 | 0 | 6 | 30.0 |
| 0 | 0 | 0 | 7 | 35.0 |
| 0 | 0 | 0 | 8 | 40.0 |
| 0 | 0 | 0 | 9 | 45.0 |
| 0 | 0 | 0 | 10 | 50.0 |
| 0 | 0 | 0 | 11 | 55.0 |

*Continued on next page...*

*Continued from previous page...*

| Number of violated restraints | | | Fraction of the ensemble | |
|---|---|---|---|---|
| PSI | PHI | Total | Count[1] | % |
| 0 | 0 | 0 | 12 | 60.0 |
| 0 | 0 | 0 | 13 | 65.0 |
| 0 | 0 | 0 | 14 | 70.0 |
| 0 | 0 | 0 | 15 | 75.0 |
| 0 | 0 | 0 | 16 | 80.0 |
| 0 | 0 | 0 | 17 | 85.0 |
| 0 | 0 | 0 | 18 | 90.0 |
| 0 | 0 | 0 | 19 | 95.0 |
| 0 | 0 | 0 | 20 | 100.0 |

[1] Number of models with violations

##### 10.3.1 Bar graph : Dihedral-angle Violation statistics for the ensemble (i)

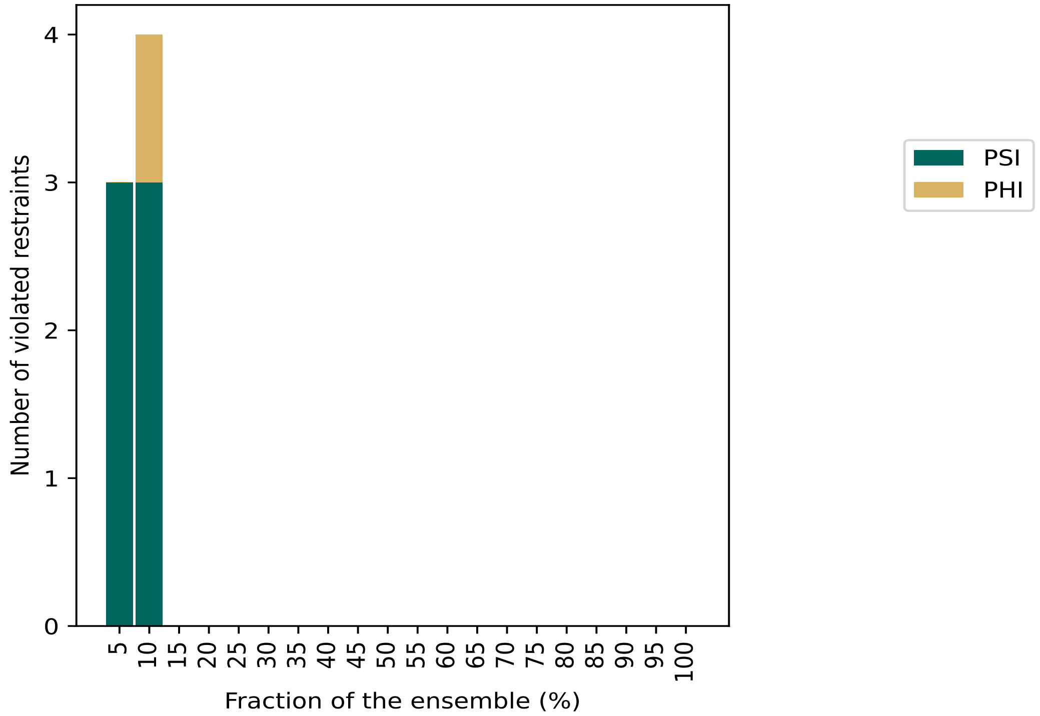


#### 10.4 Most violated dihedral-angle restraints in the ensemble (i)

##### 10.4.1 Histogram : Distribution of mean dihedral-angle violations (i)

The following histogram shows the distribution of the average value of the violation. The average is calculated for each restraint that is violated in more than one model over all the violated models

in the ensemble

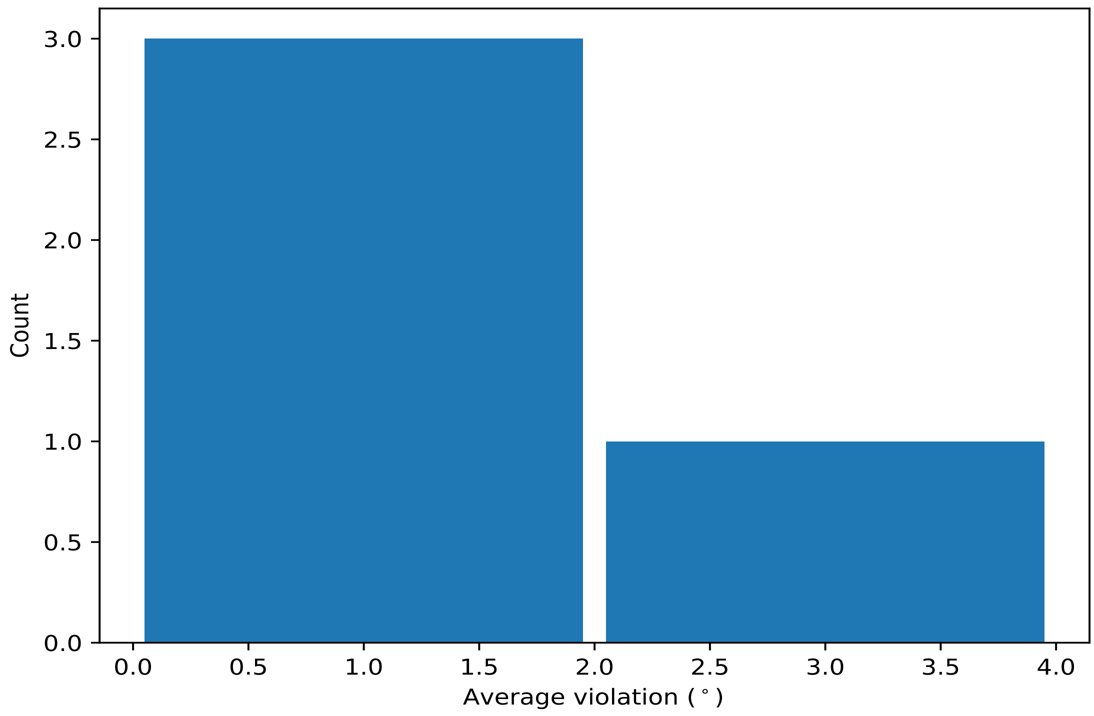


##### 10.4.2 Table: Most violated dihedral-angle restraints (i)

The following table provides the mean and the standard deviation of the violation for each restraint sorted by number of violated models and the mean value. The Key (restraint list ID, restraint ID) is the unique identifier for a given restraint.

| Key | Atom-1 | Atom-2 | Atom-3 | Atom-4 | Models[1] | Mean | SD[2] | Median |
|---|---|---|---|---|---|---|---|---|
| (1,161) | 1:A:91:GLU:N | 1:A:91:GLU:CA | 1:A:91:GLU:C | 1:A:92:VAL:N | 2 | 3.15 | 0.35 | 3.15 |
| (1,51) | 1:A:28:ASP:N | 1:A:28:ASP:CA | 1:A:28:ASP:C | 1:A:29:PHE:N | 2 | 1.5 | 0.2 | 1.5 |
| (1,129) | 1:A:68:SER:C | 1:A:69:ILE:N | 1:A:69:ILE:CA | 1:A:69:ILE:C | 2 | 1.45 | 0.15 | 1.45 |
| (1,38) | 1:A:21:TYR:N | 1:A:21:TYR:CA | 1:A:21:TYR:C | 1:A:22:LYS:N | 2 | 1.4 | 0.3 | 1.4 |

[1] Number of violated models, [2]Standard deviation, All angle values are in degree (°)

#### 10.5 All violated dihedral-angle restraints (i)

##### 10.5.1 Histogram : Distribution of violations (i)

The following histogram shows the distribution of the absolute value of the violation for all violated restraints in the ensemble.

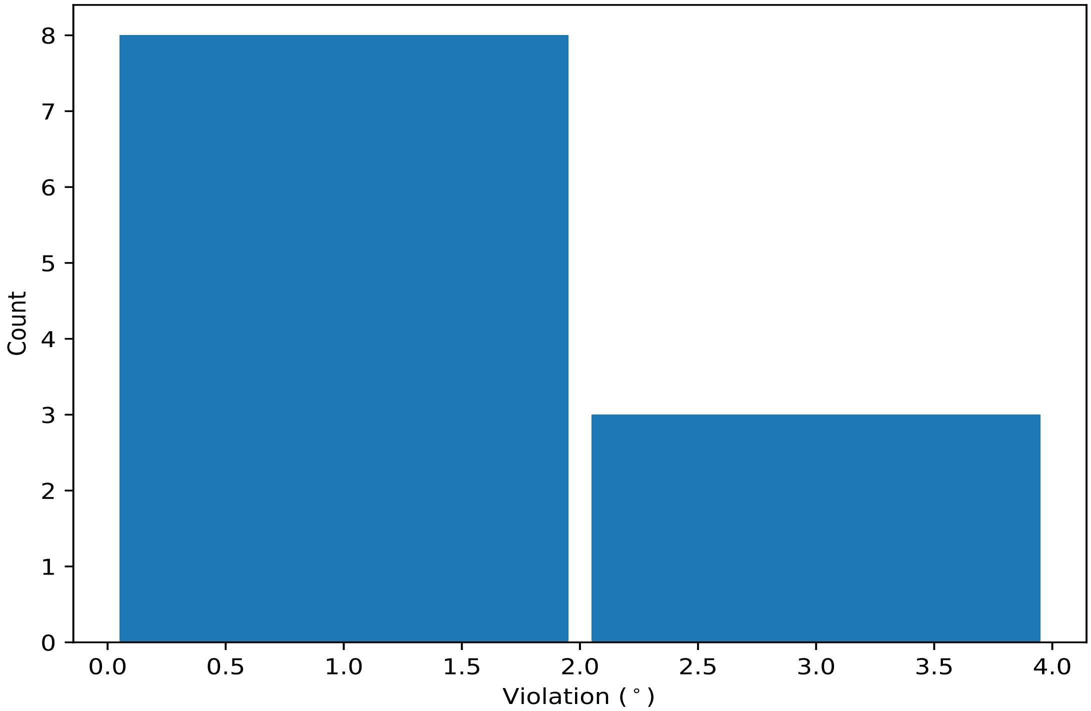


##### 10.5.2 Table: All violated dihedral-angle restraints (i)

The following table lists the absolute value of the violation for each restraint in the ensemble sorted by its value. The Key (restraint list ID, restraint ID) is the unique identifier for a given restraint.

| Key | Atom-1 | Atom-2 | Atom-3 | Atom-4 | Model ID | Violation (°) |
|---|---|---|---|---|---|---|
| (1,161) | 1:A:91:GLU:N | 1:A:91:GLU:CA | 1:A:91:GLU:C | 1:A:92:VAL:N | 6 | 3.5 |
| (1,161) | 1:A:91:GLU:N | 1:A:91:GLU:CA | 1:A:91:GLU:C | 1:A:92:VAL:N | 3 | 2.8 |
| (1,2) | 1:A:3:PHE:N | 1:A:3:PHE:CA | 1:A:3:PHE:C | 1:A:4:THR:N | 9 | 2.3 |
| (1,51) | 1:A:28:ASP:N | 1:A:28:ASP:CA | 1:A:28:ASP:C | 1:A:29:PHE:N | 3 | 1.7 |
| (1,38) | 1:A:21:TYR:N | 1:A:21:TYR:CA | 1:A:21:TYR:C | 1:A:22:LYS:N | 6 | 1.7 |
| (1,129) | 1:A:68:SER:C | 1:A:69:ILE:N | 1:A:69:ILE:CA | 1:A:69:ILE:C | 14 | 1.6 |
| (1,8) | 1:A:6:ARG:N | 1:A:6:ARG:CA | 1:A:6:ARG:C | 1:A:7:LEU:N | 19 | 1.4 |
| (1,51) | 1:A:28:ASP:N | 1:A:28:ASP:CA | 1:A:28:ASP:C | 1:A:29:PHE:N | 2 | 1.3 |
| (1,129) | 1:A:68:SER:C | 1:A:69:ILE:N | 1:A:69:ILE:CA | 1:A:69:ILE:C | 10 | 1.3 |
| (1,49) | 1:A:27:PRO:N | 1:A:27:PRO:CA | 1:A:27:PRO:C | 1:A:28:ASP:N | 13 | 1.1 |
| (1,38) | 1:A:21:TYR:N | 1:A:21:TYR:CA | 1:A:21:TYR:C | 1:A:22:LYS:N | 8 | 1.1 |
